# Dysregulation of anti-Ro60 B cell autoreactivity in systemic lupus erythematosus

**DOI:** 10.64898/2026.05.08.723865

**Authors:** Oindrila Rahaman, Carlos Castrillon, Regina Bugrovsky, Raksha Das, Midushi Ghimire, Trinh T.P Van, Martin Lin, Sabeena Y Usman, R.Toby Amoss, Aakriti Alisha Arora, Arezou Khosroshahi, Frances Eun-Hyung Lee, Ignacio Sanz

## Abstract

To understand the dysregulation of autoreactive B cells in SLE, we tracked Ro60-specific cells in seropositive (SP) and seronegative (SN) patients and healthy donors (HD), using flow cytometry and monoclonal antibodies. Consistent with permissive central tolerance, Ro60^+^ naïve B cells were present in all groups with increased anergy in HD. HD and SN SLE also had greatly decreased or absent Ro60^+^ memory and ASC, which were greatly increased in active SP SLE, thereby indicating defective distal tolerance in the latter group. Notably, Ro60 autoreactivity was strictly purged from naïve-derived extra-follicular B cells in HD and SN SLE, but expanded in SP SLE, suggesting the importance of autoreactivity censoring in this pathway. SLE clustering of the distribution of Ro60^+^ B cells identified disease heterogeneity in tolerance enforcement in SLE. Finally, we demonstrate a much higher degree of polyreactivity against other lupus antigens in SLE Ro60^+^ naïve cells, which is greatly attenuated in memory cells. Our work represents the first systematic study of antigen-specific autoreactive B cells and ASC in SLE. It enhances our understanding of human B cell tolerance and defines new approaches to measuring autoimmune activity in the course of SLE, including the assessment of immune resetting after B cell depletion therapies.

## Introduction

Systemic Lupus Erythematosus (SLE) is a complex systemic autoimmune condition characterized by widespread autoantibody production against ubiquitous self-antigens, resulting in immune-complex deposition and tissue damage^1,2^. Some SLE autoantibodies fluctuate with disease activity and may be eliminated by B cell depleting therapies (short-lived antibodies, exemplified by anti-dsDNA)^3,4^. In contrast, other autoantibodies remain consistently elevated throughout disease and after B cell therapies (long-lived autoantibodies, including anti-Ro/SS-A and other RNA-binding protein autoantibodies)^5–7^.

SLE is characterized by abnormal B cell regulation at least in part through genetically and epigenetically-determined B cell-intrinsic and extrinsic programs^8–13^. These abnormalities underpin defective tolerance against self-antigens, thereby enabling the expansion of autoreactive B cells and autoantibody-secreting cells (ASC). However, despite exponential growth in our understanding of human B cell complexity and their dysregulation in SLE, it is still unclear how disease autoreactivity is regulated across different B cell compartments. A better understanding of these central questions has been hampered by difficulties in high-throughput analysis of antigen-specific autoreactive B cells with previous studies suffering from the analysis of low cell numbers. Moreover, extant studies have focused on either polyreactivity, the study of VH4-34-associated autoreactivity or the analysis of global ANA autoreactivity without discrimination of discreet disease-related autoantigens^14–19^.

Here, we sought to characterize the distribution of Ro-60 specific B cells across all peripheral B cell compartments in human SLE. Anti-Ro60 autoantibodies, are present in large amounts in the serum of 35-70% of patients and may contribute to disease in different ways including the formation of immune complexes and the induction of type I interferon secretion^20,21^. In addition, autoreactive B cells may also play pathogenic roles through antigen presentation and activation of helper T cells. A pathogenic role is also suggested by their association with disease manifestations such as photosensitive rash, vasculitis and hypocomplementemia^22^. Notably, anti-Ro60 autoantibodies are first manifested years before the onset of clinical disease and may cross-react with the Epstein Barr Virus (EBV) antigen EBNA and commensals ^23,24^. Once generated, they remain remarkably stable even in the absence of disease activity and after B cell depletion therapies, or in the presence of anti-B cell therapies, more recently including CART cells^7^.

In this study, we addressed the regulation of anti-Ro60 autoreactivity using novel self-antigen-specific multi-parameter flow cytometry protocols and the generation of monoclonal antibodies derived from Ro60^+^ B cells at all stages of B cell differentiation. Our results quantify the frequency of Ro60^+^ cells across peripheral B cell populations and their differential regulation within these compartments in healthy donors (HD), Ro60 seronegative SLE patients (SN SLE), and Ro60 seropositive SLE patients (SP SLE), with different degrees of disease activity.

## Results

### Flow cytometry enumeration of Ro60^+^ B cells in SLE patients and healthy donors

To understand the regulation of disease-associated antigen-specific autoreactive B cells in human SLE we developed a high-throughput flow cytometric protocol to quantify anti-Ro60 B cells. To do so, tetramerized-Ro60 probes were separately labeled with APC/PE and integrated into a multidimensional flow cytometry panel that included B cell phenotypic surface markers (Supplementary Table 1). SLE samples were collected for cross-sectional analysis, with some subjects having multiple draws (Supplementary Table 2). SLE patients were first classified as Ro60 Seropositive (SP) or Ro60 Seronegative (SN) based on their historical ANA profile extracted from hospital records. Subsequently, anti-Ro60 titers were determined in our laboratory using a highly sensitive Luciferase Immunoprecipitation system (LIPS) with a dynamic range of several orders of magnitude ^25,26^. Patients were then segregated into low-disease activity (LDA), and high-disease activity (HDA) groups based on their SELENA-SLEDAI score at the time of the blood draw (Extended Data Fig. 1a). Serum anti-Ro60 LIPS titers ranged from 10^2^ - 5 x 10^6^ units in SP patients, were consistently below 10^2^ in SN patients and undetectable in HD (Fig. 1A). We further investigated the Ro60 IgG titers and anti-Ro60 B cell levels in SP patients, after splitting them cross-sectionally across different drug regimens. Most patients in LDA SP received HCQS (hydroxychloroquine) alone and/or AZA (azathioprine), while most HDA SP patients were treated with glucocorticoids (PSL: prednisolone), MMF (mycophenolate mofetil), frequently in combination with anti-BAFF therapy (BLM: belimumab) (Extended Data Fig. 2a). Given that very few patients received methotrexate (MTX) or anifrolumab (ANF), or were post-rituximab (pRTX) treatment, these drugs were excluded from further analysis. Overall, Ro60 IgG titers were comparable in all LDA SP groups. Notably, in HDA patients, anti-Ro60 titers were highest in BLM groups with lowest titers in PSL and AZA patients and intermediate titers in MMF patients. Nevertheless, the autoantibody levels were 20 times higher in patients treated with MMF in the HDA relative to LDA SP groups (Extended Data Fig. 2b). Given the lack of baseline data at the time of treatment initiation, our results cannot differentiate between drug effects and different baseline levels in the different clinical groups.

**Fig. 1.**
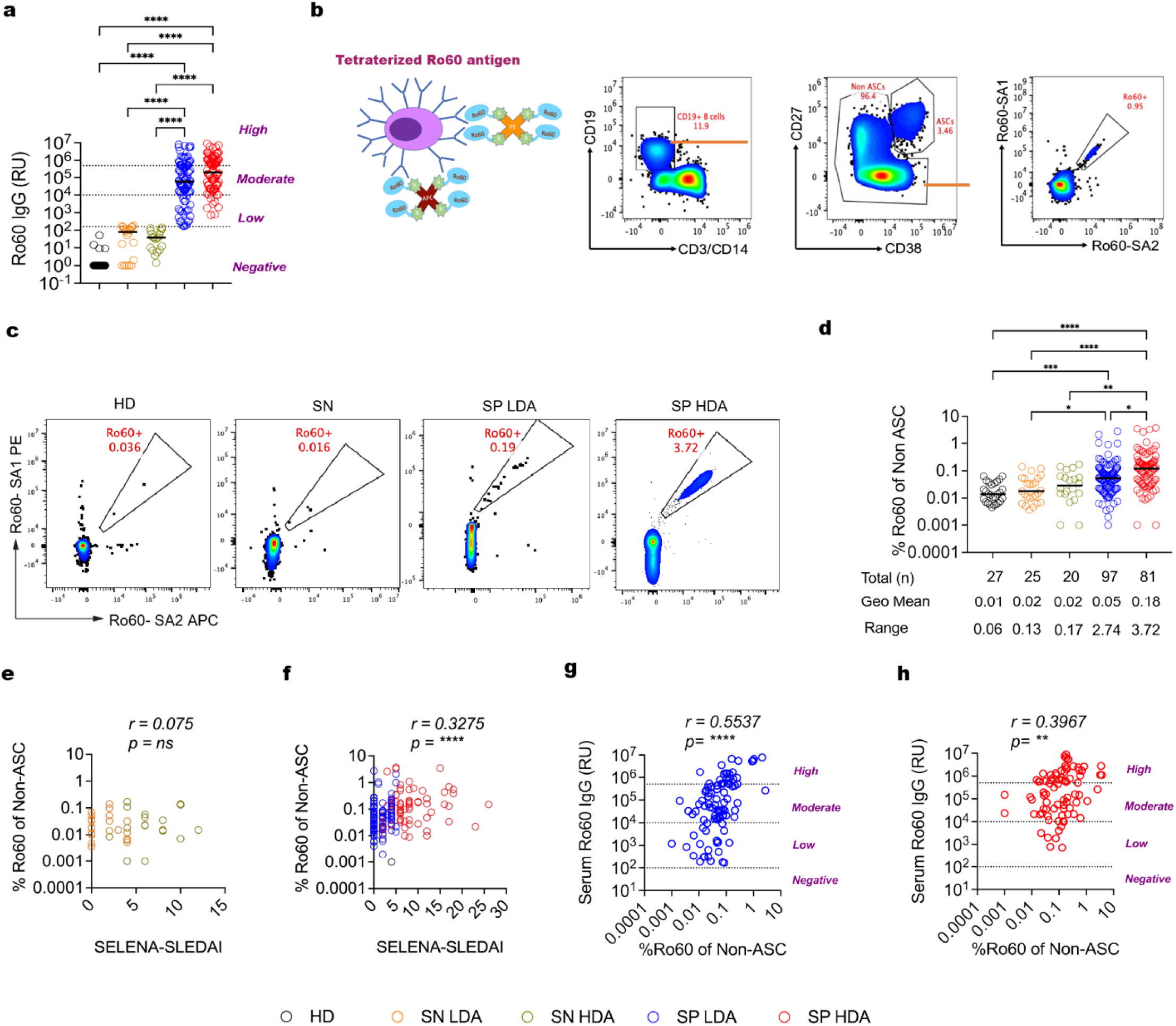
Identification and enumeration of Ro60^+^ B cells using tetrameric staining approach. **a,** Quantification of serum Ro60 IgG by LIPS assay in HD (*n*= 24), SN LDA (*n*= 20), SN HDA (*n*= 19), SP LDA (*n*= 94), and SP HDA (*n*= 81). Values below 150 RU (relative units) were considered negative. **b,** Left Panel: Cartoon representation of biotinylated Ro60 tetramers with Streptavidin-APC and Streptavidin-PE, and simultaneous binding to B cell. Right panel: Representative gating strategy for anti-Ro60 B cells. **c,** Representative anti-Ro60 flow plots as % of total B cells in HD, SN SLE, SP LDA SLE and SP HDA SLE. **d,** Scatter dot-plots of %Ro60 of B cells in HD, SN LDA, SN HDA, SP LDA and SP HDA. **e-f,** Spearman’s rank correlation of % Ro60 of B cells with SELENA-SLEDAI score in SN SLE **(e)** and SP SLE **(f)**. **g-h,** Spearman’s rank correlation of % Ro60 of B cells with serum Ro60 IgG in SP LDA **(g)** and SP HDA **(h)**. All values are plotted in log scale, values of 0 were replaced by 0.001 so that they can be plotted. Horizontal bars in scatter plots represents median, and comparison between median in different groups done by nonparametric Kruskal-Wallis Kruskal–Wallis statistical testing using Dunn’s analysis. *P < 0.05; **P < 0.01; ***P < 0.001; ****P < 0.0001.

CD19^+^ B cells were immunophenotyped into major B cell subsets defined by key cell surface markers (Supp Fig 1D). SLE patients had diminished unswitched memory (UnSwM), and increased total double negative (DN) B cells compared to HD. The activated subsets such as activated naïve (aNAV), activated SwM (aSwM), DN2, DN3 were significantly expanded in HDA SP, with a contraction of DN1 and resting subsets such as resting naïve (rNAV) and resting SwM (rSwM). Based on previous studies indicating heterogeneity in DN3 subset, we classified DN3 into the minor subset of CD38^++^ SP (single positive) and the major CD38^Neg^ subset which was similar in both HD and SLE (Extended Data Fig. 1d–l).

By flow cytometry, APC/PE double positive Ro60^+^ B cells were reliably detected in all clinical groups including HD and SN SLE patients (Fig. 1b-d and Extended Data Fig. 1j). Despite the absence of serum autoantibodies, these two groups displayed Ro60^+^ cells at mean frequencies of 0.01% to 0.02% of total CD19^+^ B cells after excluding CD27^++^ CD38^++^ antibody-secreting cells (ASC) (Fig. 1c-d). Ro60^+^ B cells were present at much higher frequencies in SP SLE, in particular in the HDA group with a mean % of 0.2, several patients scoring higher than 0.5%, and a highest value of 3.7% of all B cells. SP LDA patients were significantly higher than SN patients and HD. No differences were observed between HDA and LDA patients in the SN group (Fig. 1d). Consistently, the frequency of Ro60^+^ B cell, excluing ASC, positively correlated with SELENA-SLEDAI score in SP SLE but not in SN SLE (Fig. 1e,f). Furthermore, the Ro60^+^ B cells frequency was significantly correlated with serological Ro60 IgG titers in the LDA and HDA SP group (Fig. 1g,h). Additionally, anti-Ro60 B cells levels did not differ between the different treatment regimens (Extended Data Fig. 2c-n).

### Ro60 seropositive SLE patients display strong Ro60^+^ B cell memory

The presence of an anti-Ro60 memory compartment in SLE is inferred from the stable presence of high-affinity anti-Ro60 serum antibodies beginning years before clinical diagnosis^27,28^. Yet, direct experimental evidence and accurate measurements of such memory population is lacking. Using flow cytometry Ro60 dual staining, we first analyzed the magnitude and phenotype of anti-Ro60 CD27^+^ IgD^-^ switched memory B cells (SwM). Reflecting defective proximal peripheral tolerance, these cells were present in all groups, including HD (74%) and SN SLE (66.6%), with SP SLE displaying the highest positivity rate at 94%. In contrast, Ro60^+^ CD27^+^ IgD^+^ unswitched memory cells (UnSwM) were detectable in a smaller fraction of subjects across all groups including HD. High levels of Ro60^+^ B cells were observed in both the UnSwM and SwM compartments of SP SLE patients with frequencies as high as 6.47% and 7.76%, respectively. Nevertheless, the median frequency for SwM was significantly higher in SP HDA relative to SP LDA (Fig. 2a,b and Extended Data Fig. 6a,b). Presence of Ro60^+^ UnSwM was also associated with higher disease activity (Extended Data Fig. 1b,c). No significant differences were detected between SN patients and HD whether for either memory subsets. Importantly, Ro60^+^ memory B cell frequencies far exceeded steady-state memory for tetanus toxoid, which rarely surpassed 0.5% in a subset of the same SP SLE patients (Extended Data Fig. 3a-c). A strong positive correlation of Ro60^+^ SwM levels with the serum levels of anti-Ro60 IgG antibodies was found in both, the LDA and HDA SP groups (Fig. 2c).

**Fig. 2.**
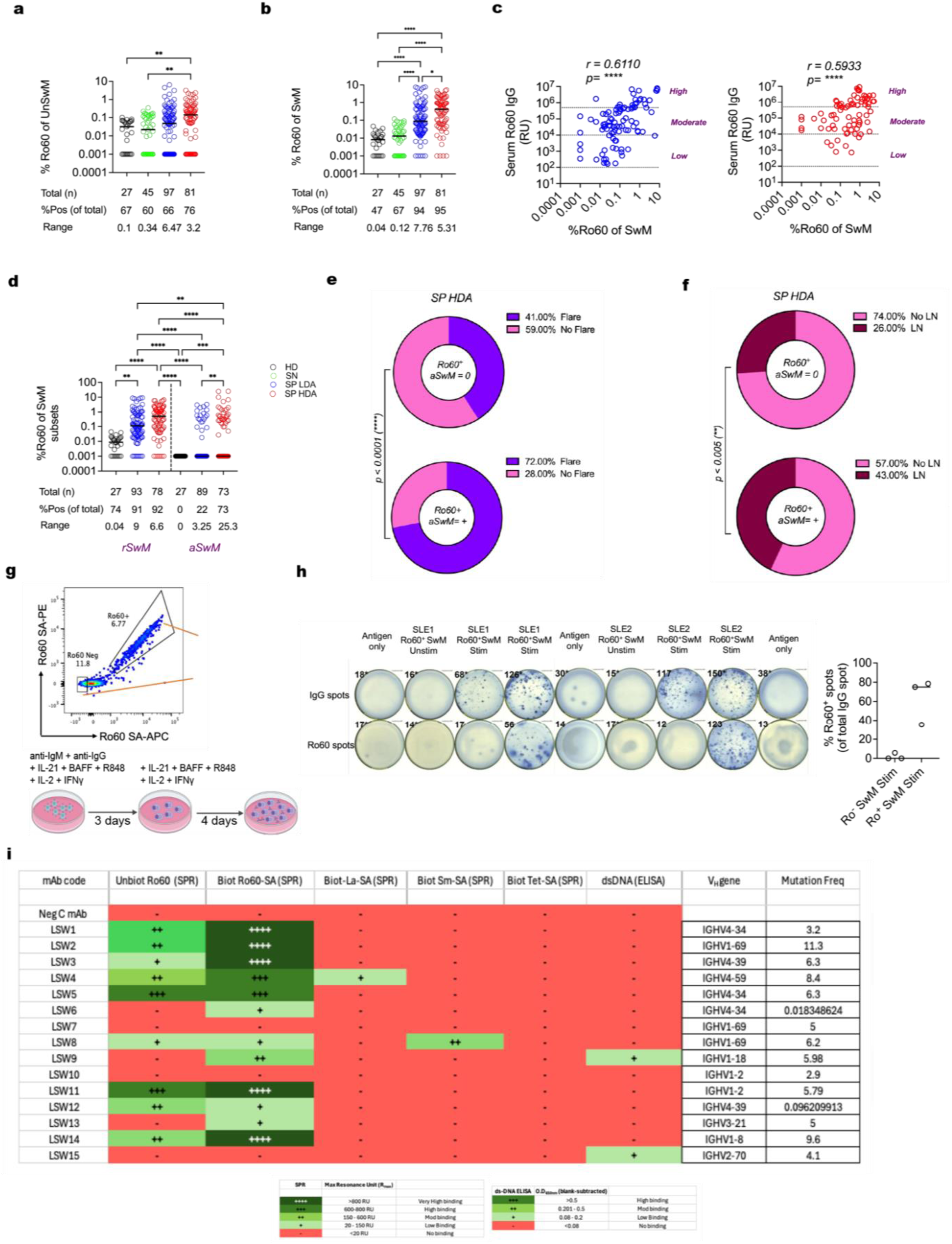
Prominent features of Ro60^+^ memory in SP SLE and their association with disease severity. Scatter dot-plots of % Ro60 of UnSwM **(a)**, SwM **(b)** in different groups. **c**, Spearman’s rank correlation of % Ro60 of SwM with serum Ro60 IgG in SP LDA **(Left panel)** and SP HDA **(Right panel)**. **d,** Scatter dot-plots of % Ro60 of rSwM (CD21^+^ CD11c^+/-^) and aSwM (CD21^-^ CD11c^+^) in different groups. **e,f,** Donut plots of proportion of HDA SP samples which are flaring vs not flaring **(e)**, or which have active LN (lupus nephritis) vs no LN **(f)**. **g,** Cartoon representation of ex-vivo culture to differentiate Ro60^+^ and Ro60^-^ SwM into ASC. **h,** Left panel: Representative anti-Ro60 and anti-IgG ELISPLOT of unstimulated Ro60^-^ SwM, stimulated Ro60^-^ SwM and Ro60^+^ SwM. Cells seeded per well for SLE1 was 650 cells, and for SLE2 was 1750 cells. Right panel: Scatter dot plot of % Ro60-specific of total IgG ASC in stimulated Ro60^-^ SwM and Ro60^+^ SwM (normalized over unstimulated Ro60^-^ SwM) from 3 subjects. **i**, Heatmap of Rmax (Resonance Unit max) plots from SPR binding studies (unbiotinylated Ro60, biotinylated Ro60-SA tetramer, biotinylated La-SA tetramer, biotinylated Sm-SA tetramer, and biotinylated Tet-SA tetramer) or O.D650nm(Blank subtracted) from anti-ds-DNA ELISA for 15 monoclonals derived from Ro60^+^ SwM (LSW1 to LSW15). For scatter plot and correlation plots, all values are plotted in log scale, values of 0 were replaced by 0.001 so that they can be visualized. Horizontal bars in scatter plots represents median, and comparison between median in different groups done by nonparametric Kruskal-Wallis Kruskal–Wallis statistical testing using Dunn’s analysis. Donut plots were compared by Chi square test.

Next, we measured the relative frequency of Ro60^+^ cells within the activated fraction of the SwM compartment. Notably, Ro60^+^ activated SwM cells (aSwM, CD21^-^ CD11c^+^), were undetectable in HD, absent in a large fraction of SP HDA patients (52%), and a large majority of SP LDA (78%). In contrast, the vast majority of SP patients in both activity groups displayed detectable numbers of resting Ro60^+^ SwM cells (Fig. 2d, Extended Data Fig. 6c). Nevertheless, we observed that within the smaller group of HDA SP patients, those with detectable anti-Ro60 aSwM cells, had significantly higher SELENA-SLEDAI scores, a significantly higher rate of flares and more prevalence of active lupus nephritis (Fig. 2e,f and Extended Data Fig. 1b,c). Importantly, the levels of anti-Ro60 rSwM and aSwM were not different across treatment groups including patients treated with BLM (Extended Data Fig. 2h-l). Combined, these results support the counterintuitive possibility that activation of autoreactive memory may not be necessary for disease activation. Longitudinal analysis of LDA patients may be required to formally assess the significance of activation of the anti-Ro60 memory response in disease manifestations (including lupus nephritis) and flares.

To validate the ability of flow staining to detect Ro60-specific B cells, we sorted Ro60^+^ SwM B cells from SP patients and differentiated them in-vitro, under TH1-like stimulation conditions, into antibody secreting cells (ASC) (Fig. 2g). As indicated by ELISPOT assays, anti-Ro60 ASC developed in Ro60^+^ SwM wells and but not in Ro60^-^ SwM wells, although IgG spots were present in both (Fig. 2h). To further validate the specificity of Ro60 staining, we generated monoclonal antibodies (mAbs) from sorted Ro60^+^ SwM B cells and assessed their binding to Ro60 antigen by surface plasmon resonance (SPR). The monoclonal antibodies selected for testing represented a range of different VH gene usage and different degrees of somatic hypermutation (Extended Data Fig. 4a-e and Supplementary Table 3). Binding to unbiotinylated monomeric Ro60 was detected by SPR in 60% of the mAbs tested and in 80% when using biotinylated Ro60-SA tetramers to increase antigen density. In addition, the binding magnitude (Rmax) was consistently higher with biotinylated Ro60-SA tetramers compared to monomeric Ro60. Excluding the contribution of non-specific mAb binding to biotin-streptavidin epitopes, there was no binding to biotinylated tetanus-SA tetramer. When tested against other lupus autoantigens (Smith, dsDNA and La), polyreactivity was absent in most mAbs, which did not react either against biotinylated tetanus toxoid (Fig. 2i).

### Unmutated naïve B cells contain anti-Ro60 autoreactivity and are dysregulated in active SLE

The presence of autoreactive naïve B cells indicates failure of central tolerance. Hence, we explored the presence and properties of anti-Ro60 autoreactivity in this compartment. Resting naïve (rNAV) B cells contained detectable levels of Ro60^+^ B cells in all HD and in a large majority of patients in all SLE groups (Fig. 3a and Extended Data Fig. 6d). Of interest, almost all SLE patients with undetectable Ro60^+^ B cells (5-16%, in the different groups), were treated with Belimumab, a finding consistent with the ability of BAFF blockade to regulate the survival of autoreactive naïve B cells ^29^ (Extended Data Fig. 2e). Notably, Ro60^+^ resting naïve cells displayed an IgD^+^ IgM^-^ phenotype in the majority of HD at a much higher frequency than SLE patients, which is consistent with anergy which is the main mechanism of early peripheral tolerance^30,31^ (Fig. 3b).

**Fig. 3.**
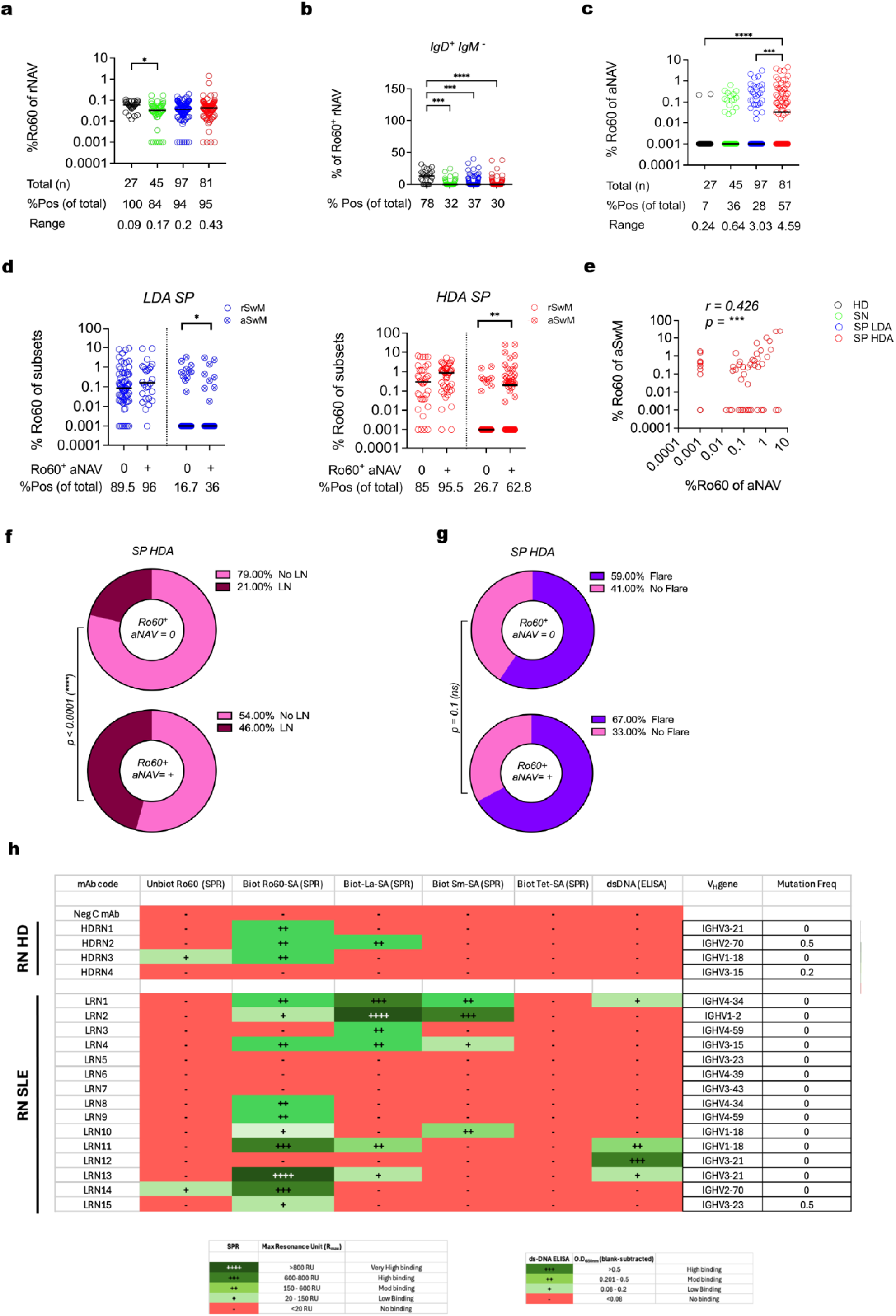
Unmutated naïve B cell compartment harbors Ro60^+^ autoreactivity. **a-c,** Scatter dot-plots of % Ro60 of rNAV **(a),** %IgD^+^ IgM^-^ of Ro60^+^ rNAV **(b)** and % Ro60 of aNAV **(c)** in different groups. **d,** Scatter dot-plot of % Ro60 of rSwM and aSwM in LDA SP (Left panel) and HDA SP (Right panel) patients with (denoted by +) and without (denoted by 0) detectable Ro60^+^ aNAV. **e,** Spearman’s rank correlation of % Ro60 of aSwM and %Ro60 of aNAV in HDA SP. **f,g,** Donut plots of proportion of HDA SP samples which have active LN (lupus nephritis) vs no LN **(f)**, and which are flaring vs not flaring **(g)**. **h,** Heatmap of Rmax (Resonance Unit max) plots from SPR binding studies (unbiotinylated Ro60, biotinylated Ro60-SA tetramer, biotinylated La-SA tetramer, biotinylated Sm-SA tetramer, and biotinylated Tet-SA tetramer) or O.D650nm(Blank subtracted) from anti-ds-DNA ELISA of anti-Ro60 resting naïve derived monoclonals. 4 monoclonals were made from HD Ro60^+^ rNAV (HDRN1 to HDRN4) and 15 monoclonals were made from SP SLE Ro60^+^ rNAV (LRN1 to LRN15). For scatter plot and correlation plots, all values are plotted in log scale, values of 0 were replaced by 0.001 so that they can be visualized. Horizontal bars in scatter plots represents median, and comparison between median in different groups done by nonparametric Kruskal-Wallis Kruskal–Wallis statistical testing using Dunn’s analysis. Donut plots were compared by Chi square test. *P < 0.05; **P < 0.01; ***P < 0.001; ****P < 0.0001.

Despite their presence in the rNAV compartment, Ro60^+^ aNAV B cells were undetectable in HD, but present at high frequencies in over 50% of SP SLE patients with high disease activity, and in a smaller fraction of SN patients (36.3%), and of SP patients with LDA (27.8%) (Fig. 3c and Extended Data Fig. 6e). Presence of Ro60^+^ aNAV cells was associated with a significant increase in Ro60^+^ aSwM cells, which was much more pronounced in HDA patients (Fig. 3d). Consistently, Ro60^+^ aNAV and showed a positive correlation with Ro60^+^ aSwM levels in HDA SP (Fig. 3e). Clinically, an anti-Ro60 activated naïve compartment was associated with a higher rate of active LN in HDA patients but not with differences in disease flares (Fig. 3f,g).

As for memory cells, the specificity of Ro60^+^ naïve B cells detected by flow cytometry, was validated through the analysis of mAbs generated from Ro60^+^ rNAV cells of SP SLE and one healthy donor (Extended Data Fig. 4a-e). Binding to biotinylated Ro60-SA tetramer was detected in 66.7% and 75% of SLE and HD rNAV-derived monoclonals, respectively. In contrast to memory B cells, anti-Ro60^+^ lupus resting naïve B cells displayed preferential binding to tetrameric antigen while generally failing to react with unbiotinylated monomeric Ro60, a pattern consistent with the low affinity binding expected in unselected naïve B cells. Also, in contrast to mAb derived from Ro60^+^ memory cells, mAbs from SLE naïve cells displayed a substantial degree of polyreactivity against other lupus-associated autoantigens prominently including La, dsDNA and Smith. Polyreactivity was less pronounced in mAb derived from Ro60^+^ healthy naïve cells. No reactivity was detected against biotinylated tetanus antigen, indicating the mAbs did not bind to biotin and the preferential binding of anti-Ro60 naïve B cells to self-antigens (Fig. 3h and Supplementary Table 3).

### Ro60 autoreactivity present at high levels in different subsets of double negative B cells in both LDA and HDA disease

IgD/CD27 double negative (DN) B cells, which are greatly expanded in active SLE, are heterogenous and can be classified in DN1, DN2 and DN3 B cells based on differential expression of CD21 and CD11c. DN1 (CD21^+^ CD11c^-^), are thought to represent early memory populations, while DN2 (CD21^-^ CD11c^++^), represent naïve-derived cells epigenetically poise to differentiate into plasmablast. DN2 cells are enriched in SLE-related autoreactivity and expanded in Black patients with lupus nephritis^25,26^. Both subsets contribute durable memory in response to vaccination^32,33^. Finally, DN3 cells (CD21^-^ CD11c^-^), are considered pre-plasmablasts that may infiltrate end organs in COVID-19 and IgG4- related disease, and are also expanded in active SLE patients^34–38^. Ro60^+^ cells were present in all DN subsets in the majority of SP SLE patients in both LDA and HDA groups (60-89.7%, respectively), at frequencies reaching as high as 5-10% of the different DN populations with highest positivity in SP HDA patients (Fig. 4a-c and Extended Data Fig. 6 f-h). Notably, Ro60 DN1 positivity and levels were significantly lower than rSwM (Fig. 2b and Extended Data Fig. 5c-e), thereby suggesting uncoupling of CD27^+^ and CD27^-^ memory compartments.

**Fig. 4.**
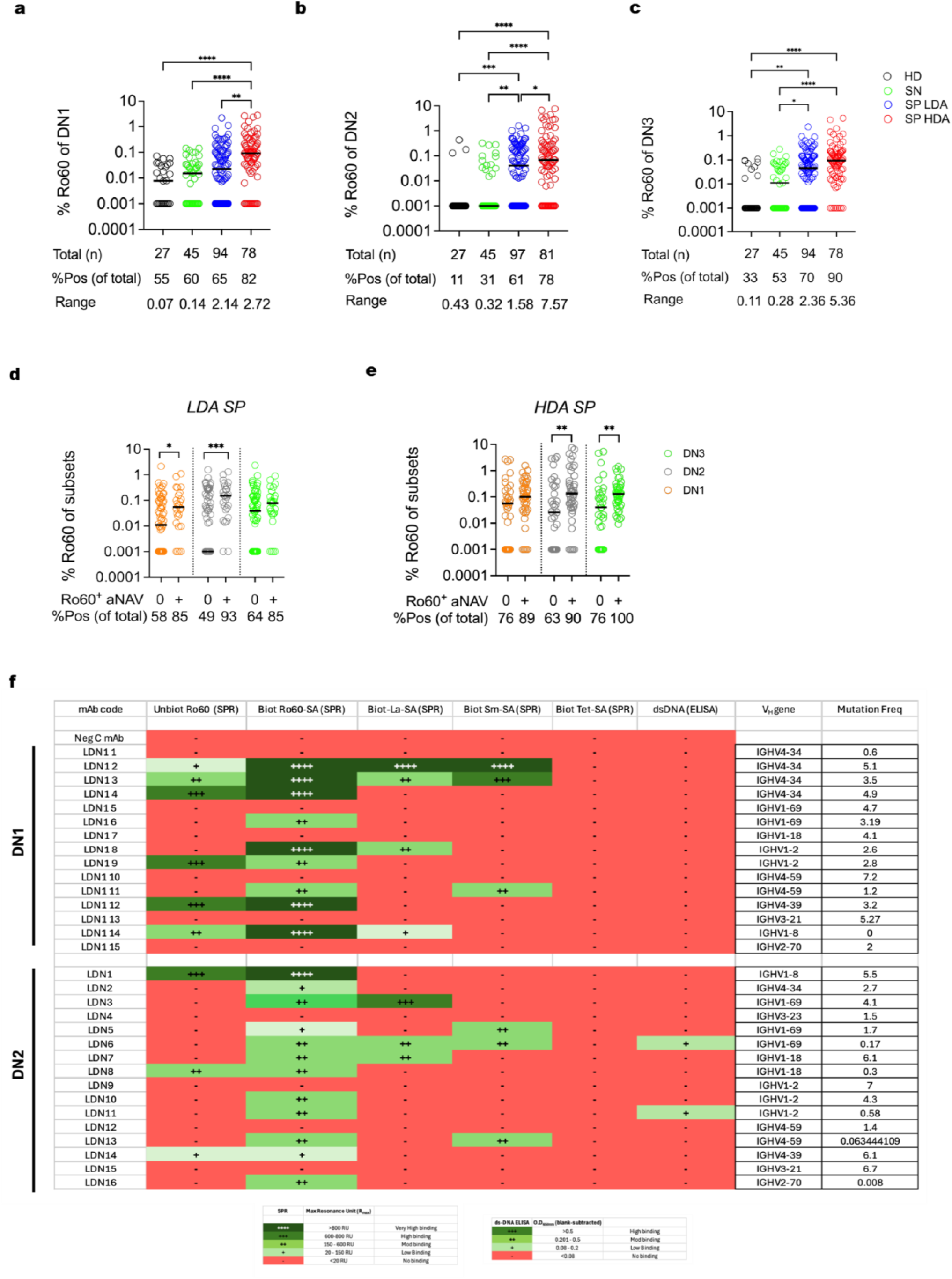
Elevated Ro60^+^ autoreactivity across DN B cell subsets in SP SLE. **a-c)** Scatter dot-plots of % Ro60 of DN1 **(a),** DN2 **(b)** and DN3 **(c)** in different groups. **d,** Scatter dot-plot of % Ro60 of DN1, DN2, DN3 in LDA SP (Left panel) and HDA SP (Right panel) patients with (denoted by +) and without (denoted by 0) detectable Ro60^+^ aNAV. **e,** Heatmap of Rmax (Resonance Unit max) plots from SPR binding studies (unbiotinylated Ro60, biotinylated Ro60-SA tetramer, biotinylated La-SA tetramer, biotinylated Sm-SA tetramer, and biotinylated Tet-SA tetramer) or O.D650nm(Blank subtracted) from anti-ds-DNA ELISA of 15 monoclonals made from anti-Ro60 DN1 (LDN1 1 to LDN1 15) and 16 monoclonals from anti-Ro60 DN2 (LDN1 to LDN16). For scatter plot and correlation plots, all values are plotted in log scale, values of 0 were replaced by 0.001 so that they can be visualized. Horizontal bars in scatter plots represents median, and comparison between median in different groups done by nonparametric Kruskal-Wallis Kruskal–Wallis statistical testing using Dunn’s analysis. *P < 0.05; **P < 0.01; ***P < 0.001; ****P < 0.0001.

Ro60^+^ cells were almost absent in HD DN2, and present at low levels in 30% of SN SLE, with higher levels present in SP SLE, with highest levels found in HDA SP subjects. These findings are consistent with strict censoring of Ro60^+^ cells within the effector DN2 compartment in HD, faulty censoring in SP SLE and expansion in HDA SLE. A very similar pattern was observed for DN3 cells. The combined DN2/DN3 findings are consistent with active differentiation of Ro60^+^ B cells into plasmablasts, even in SP patients with LDA (Extended Data Fig. 5a-e). Nevertheless, this process was particularly prominent in SP patients with HDA, a group in which the presence of Ro60^+^ DN2/DN3 cells was significantly associated with higher SLEDAI scores (Extended Data Fig. 1b,c and Extended Data Fig. 5e,f). Notably, Ro60^+^ DN2^High^ patients in HDA group had higher SELENA-SLEDAI scores, higher serum anti-Ro60 IgG and a significantly higher incidence of active lupus nephritis, than Ro60^+^ DN2^low^ patients (Extended Data Fig. 7a-d). Consistent with the naïve-derived model, the patients with anti-Ro60 aNAV B cells also had significantly more Ro60^+^ DN2 in both LDA and HDA SP SLE (Fig 4D).

As for other populations, we used mAbs to characterize the autoreactivity of Ro60^+^ DN cells. The majority of mAb derived from Ro60^+^ DN1 and DN2 cells (60% and 75%, respectively), bound to biotinylated Ro60-SA, thereby validating the specificity of Ro60 binding by flow cytometry. However, a greater proportion of DN1-derived mAbs (40%) were able to bind unbiotinylated monomeric Ro60 compared to DN2-derived mAbs (19%), and DN1-derived mAbs had higher overall binding to biotinylated Ro60. The lower binding of DN2 derived antibodies was associated with a lower degree of somatic hypermutation. Notably, in both groups, 33% of antibodies demonstrated significant polyreactivity against other RNA-binding proteins, La and/or Smith. In addition, two anti-Ro60 antibodies from DN2 cells also demonstrated low binding to dsDNA, in one case together with Smith binding, a behavior previously identified in lpr mice ^39,40^ (Fig. 4e, Extended Data Fig. 4a-e and Supplementary Table 3). The lower SHM, lower Ro60 binding and widespread poly-reactivity of DN2 antibodies, would be consistent with a naïve-derived origin and extrafollicular differentiation of DN2 cells ^25^. In turn, DN1 antibodies displayed intermediate features between DN2 and SwM antibodies, consistent with an early GC derivation previously proposed for these cells^12^.

### Analysis of anti-Ro60 antibody-secreting cells and precursor responses using intracellular staining

While SLE is characterized by the peripheral expansion of ASC in correlation with disease activity, their degree of autoreactivity is not well understood as only limited ANA-reactivity analysis has been conducted using surface staining using flow cytometry. However, this approach is limited to the detection of a small fraction of early plasmablasts that still retain expression of surface BCR. Yet, we have shown that the majority of blood ASC are more mature plasmablasts and plasma cells that lack surface BCR^13^. We have also reported that DN2 cells are epigenetically primed to differentiate into ASC and that, consistent with that feature, express low levels of surface BCR ^12^. In addition, DN3 cells (also increased in active SLE), have been proposed to represent pre-plasmablasts based on both phenotypic and transcriptional analysis^41^. However, the expression of surface BCR has not been ascertained in DN3 cells. Due to these considerations, an accurate analysis of the frequency of antigen-specific autoreactive ASC and their precursors may only be obtained through the use of intracellular staining (ICS), to compare the intensity of surface and cytoplasmic binding to labeled antigen. ICS of CD19^+^ IgD^-^ CD38^++^ CD27^++^ ASC, revealed that the majority of HDA SP patients (89.9%), had detectable circulating Ro60^+^ ASC relative to LDA SP patients (71.6%), with higher % of Ro60^+^ ASC in HDA (Fig. 5a-c and Extended Data Fig. 6l). Consistent with our previous data using an ELISPOT approach^42^, this approach revealed that the frequency of these cells may reach 9% of all circulating ASC and that their frequency positively correlates with serum Ro60 IgG levels in both activity groups. Interestingly, there were some SP patients without any detectable Ro60^+^ ASC. Hence, while the stable levels of serum anti-Ro60 IgG antibodies, even in the presence of profound B cells and CD19^+^ ASC depletion, indicates the presence of a Ro60^+^ CD19- long-lived plasma cell bone marrow compartment ^7^. Our data provide the first evidence for the ongoing generation of Ro60^+^ ASC even at the time of low disease activity with increased generation during phases of high disease activity. Ro60^+^ ASC were absent in HC and present at low levels in about 40% of SN SLE.

**Fig. 5.**
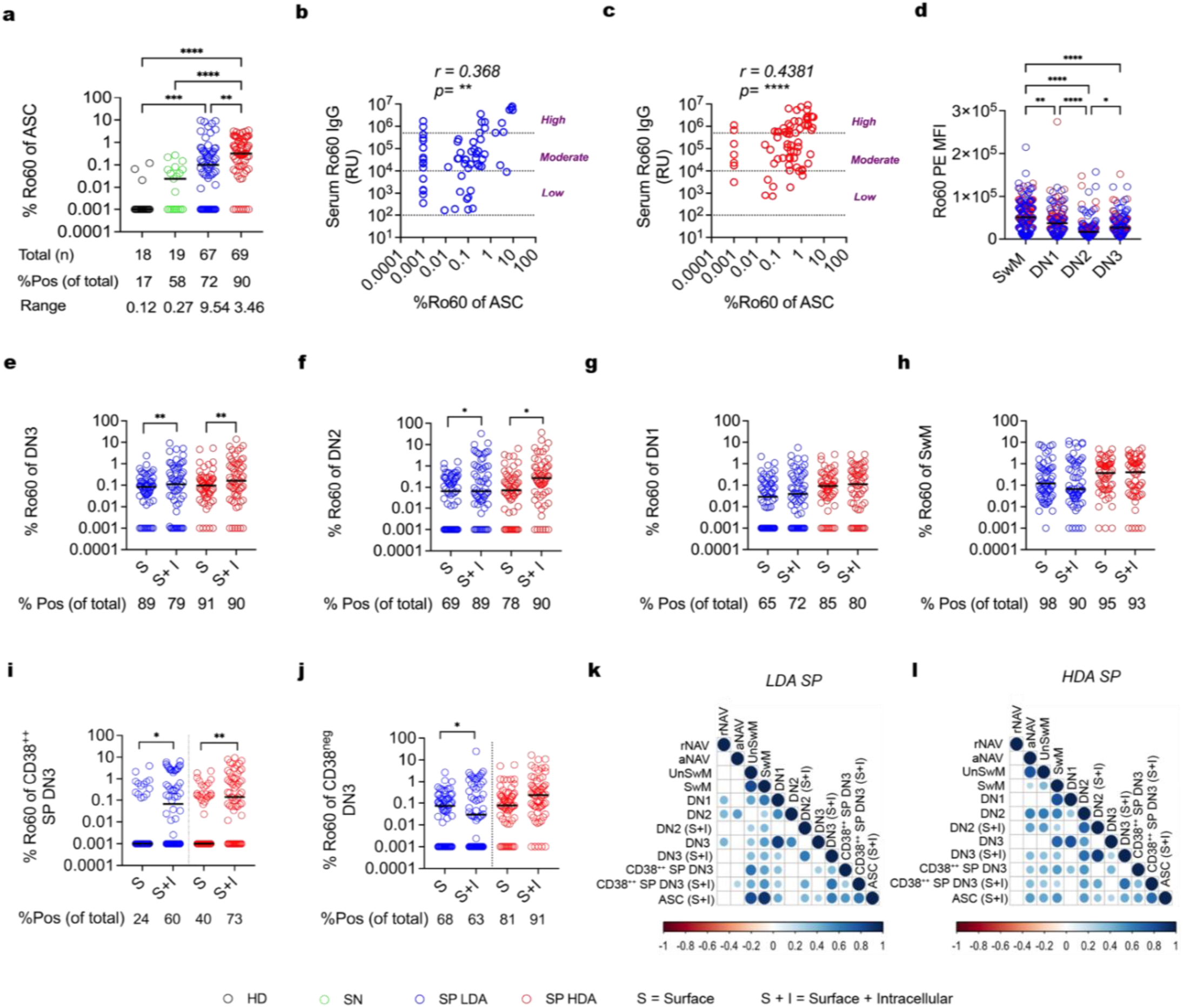
Robust Ro60-specific ASC generation in LDA SP SLE, and expansion in both groups of HDA SP SLE. **a,** Scatter dot-plots of % Ro60 of ASC (antibody-secreting cells) by intracellular staining in different groups. **b, c,** Spearman’s correlation of % Ro60 in ASC with serum Ro60 IgG titers in LDA SP **(b)** and HDA SP **(b)**. D) Scatter dot-plots of anti-Ro60 PE MFI in SwM, DN1, DN2, and DN3. **e-j,** Scatter dot-plots for comparison of % Ro60^+^ frequency on surface (S) versus intracellular (S+I) in DN3 **(e)**, DN2 **(f)**, DN1 **(g)**, SwM **(h)**, CD38^++^ SP DN3 **(i)** and CD38^Neg^ DN3 **(j)**. **k,l,** Correlograms of Spearman correlation of %Ro60 of different subsets in LDA SP **(k)** and HDA SP **(l)**. All values are plotted in log scale, values of 0 were replaced by 0.001 so that they can be plotted. Horizontal bars in scatter plots represents median, and comparison between median in different groups done by nonparametric Kruskal–Wallis statistical testing using Dunn’s analysis for **a-d**. Median was compared pairwise by Wilcoxon signed-rank test for **e-j**. *P < 0.05; **P < 0.01; ***P < 0.001; ****P < 0.0001.

Among non-ASC, we found a gradient of MFI for surface Ro60 staining with highest levels in SwM, followed by DN1 cells, with lowest levels in DN2 and DN3, consistent with differences in surface BCR previously reported^25^ (Fig. 5d). ICS was then performed to determine accumulation of intracellular BCRs, in order to optimize the detection of Ro60^+^ cells. A significant increase in Ro60^+^ frequencies in ICS relative to surface staining was evident in DN3 and DN2 only with no differences observed for SwM and DN1 B cells (Fig. 5e-h). Notably, ICS detected Ro60^+^ DN2 in about 90% of SP patients irrespective of disease activity with frequencies as high as 30% of DN2 cells. These data first document that these B cells display a central feature of an ASC lineage that is not shared by SwM or DN1 B cells. Importantly, the % of Ro60^+^ DN2 and DN3 cells that acquire this ASC feature increases with disease activity. Also of note, intracellular Ro60^+^ BCR accumulation in DN3 cells was highly significant in the CD38^++^ SP subset for SP SLE while being absent in HD and SN, presumably representing the pre-plasmablast population previously reported by others. In contrast, the larger CD38^neg^ fraction, representing the majority of DN3 cells, did not share this feature (Fig. 5i,j and Extended Data Fig. 5j).

Finally, we investigated the correlation between Ro60^+^ ASC and other Ro60^+^ subsets. We generated a correlation matrix of Ro60^+^ cells in all subsets. Ro60^+^ ASC were strongly correlated with anti-Ro60^+^ SwM, UnSwM, and moderately correlated with intracellular DN3 frequencies, in particular in the CD38^++^ DN3 fraction. In HDA, while ASC correlations with memory cells were weaker, ASC correlations with aNAV and DN2 cells became apparent. It is interesting to note that none of the drugs were effective in reducing Ro60^+^ ASC, which was similar to our finding of similar Ro60 levels in pre-plasmablast DN3 and similar Ro60 IgG titers across all drug groups (Extended Data Fig. 2d,l-n).

## Discussion

Mouse studies, largely performed in animals transgenic for different antigens, have established the existence of multiple tolerance checkpoints for the censoring of antigen-specific autoreactive B cells in order to ultimately avoid the generation of autoantibodies and autoimmune disease. Combined, these studies have demonstrated the presence of central checkpoints during bone marrow B cell development that prevent the accumulation of peripheral mature naïve B cells, through either clonal deletion, or receptor editing^43–47^. Additional checkpoints have been identified that prevent the activation, survival and/or differentiation into ASC, of autoreactive B cells that escape central tolerance (peripheral tolerance)^48^. In turn, peripheral tolerance may be enforced early during activation of naïve B cells (through anergy, follicular exclusion and decreased survival), or during differentiation and selection in the germinal centers (GC; activation induced cell death, FAS-mediated mechanisms and receptor revision among other mechanisms)^49^. Finally, late peripheral tolerance may operate on memory B cells to prevent their final differentiation into ASC through mechanisms not fully understood^50^. Human B cell tolerance is in contrast, poorly understood with limited studies of antigen-specific cells and other experimental limitations. Extant studies have concentrated on measurements of global ANA and polyreactivity against non-disease relevant antigens, using limited numbers of patients and monoclonal antibodies, to propose the presence of both central and peripheral checkpoints (both early and late). These mechanisms would progressively censor autoreactive B cells to low but significant numbers of memory cells and additional censoring into the bone marrow ASC compartment. In these studies, these tolerogenic checkpoints were faulty in SLE^17,51–53^. In contrast, larger studies using flow cytometry to study global ANA autoreactivity, have proposed the preservation of central and peripheral checkpoints in SLE relative to HD. In this model, the increased autoreactivity characteristic of SLE would be accounted for by polyclonal B cell activation leading to accumulation of autoreactive ASC^18,19^. Finally, our group and others have demonstrated that the intrinsic autoreactivity of B cells expressing VH4-34-encoded B cell receptors is censored in HD through GC censoring and redemption through somatic hypermutation^16,17,54^. Of significant interest, it is generally accepted that the differentiation of autoreactive naïve B cells into ASC through an extrafollicular pathway of high pathogenic potential both in SLE and viral infections, may not be censored through specific tolerogenic checkpoints. Instead, this outcome would be prevented by the derivation of these cells to a GC pathway with operational checkpoints^55,56^.

In the present study, we sought to understand human B cell tolerance and its breakdown in SLE through the analysis of the fate of cells that recognize the SLE-associated self-antigen Ro60. This autoreactivity is also informative since anti-Ro autoantibodies are the first to be identified in SLE and are present years before disease onset, are associated with clinical manifestations and as immune complexes, are powerful inducers of type I interferon^21,22,27^. Experimentally, our approach benefits from several advantages including the use of a single disease-associated antigen rather than ANA preparations containing large numbers of self-antigens, the high-throughput afforded by the use of flow cytometry and the binary nature of SLE with respect to the presence of anti-Ro60 serum autoantibodies. The latter feature enabled us to compare SLE patients with effective tolerance (Ro60 seronegative patients; SN), in contrast to patients with defective tolerance and serum autoantibodies (Ro60 seropositive patients; SP). This is in contrast to the virtually universal seropositivity for ANA autoantibodies analyzed in other studies. In this study, we determined the frequency of autoreactive Ro60^+^ B cells along the differentiation trajectories of naïve B cells through extra-follicular (EF), and memory pathways and mapped defective tolerance checkpoints in SLE patients with effective or faulty global tolerance as indicated by the absence or presence, respectively, of serum anti-Ro60 autoantibodies. Our data demonstrate a universal defect in central tolerance leading to the presence of similar levels of Ro60^+^ naïve B cells in HD and SLE patients, whether SP or SN for anti-Ro60 autoantibodies, thereby highlighting the need for effective peripheral checkpoints downstream from resting naïve B cells. Indeed, an operational proximal tolerance checkpoint in HD is indicated by both, the absence of Ro60^+^ aNAV B cells and a significant decline in the frequency of Ro60^+^ cells between rNAV and rSwM/DN1 memory cells. This negative regulation could be accounted for at least in part, by anergy of resting naïve cells, which appears to be defective in all SLE groups. Interestingly, detectable levels of both Ro60^+^ aNAV and SwM cells, were present in a fraction of SN patients (cluster C4 and C5) suggesting the participation of late peripheral checkpoints to prevent the generation of Ro60^+^ ASC and autoantibodies in these patients (Extended Data Fig. 5a,b,e). Notably, SP patients with undetectable Ro60^+^ naïve B cells were accumulated in the BLM group, a finding consistent with the ability of BAFF blockade to inhibit the survival of autoreactive naïve B cells and the dependence of anergic B cells on BAFF for survival^8,29,57^

Our study also provides evidence to suggest that the differentiation of autoreactive Ro60^+^ naïve B cells in HD may be differentially censored along extrafollicular (EF; activated naïve, DN2 and DN3 cells), and GC-dependent memory pathways (CD27^+^ switched memory and DN1 cells), with strict censoring for EF differentiation yet, permissive differentiation into resting, central memory cells. This differential regulation also appears to prevent the activation of Ro60^+^ naïve and memory cells in HD but not in SLE, which displayed different levels of tolerance breakdown largely associated with serological status and to a significant extent with disease activity. Thus, while Ro60^+^ switched memory cells were present in all clinical groups, only SP patients displayed higher memory values relative to naïve values with highest SwM/rNAV ratios in HDA patients. Together with parallel expansions of Ro60^+^ ASC, these observations are consistent with tolerance breakdown at late peripheral checkpoints of higher magnitude of expansion in patients with HDA. Notably however, a significant majority of SP patients (cluster C10) did not display activation of the Ro60^+^ cells in the SwM compartment even in the presence of HDA and expansion of these cells in the aNAV compartment (Extended Data Fig. 5c-e,h). Whether this observation also applies to other autoreactivities, remains to be determined by other antigen-specific studies. These data lend credence to the model previously put forth by our lab where the recruitment of newly activated naïve B cells would be a prominent feature of active SLE even in the presence of abundant autoreactive memory, with aNAV cells contributing a large fraction of newly generated ASC ^25,42^. Further, the data support the notion that activation of Ro60 memory is neither a required driver nor a universal consequence of active SLE. It should be noted however, that HDA patients with this feature were significantly more likely to have active lupus nephritis and/or be experiencing a flare at the time of the study. Longitudinal analysis of LDA patients will be required to formally assess the significance of activation of the anti-Ro60 memory response in disease manifestations (such as lupus nephritis) and flares.

We and others have shown that general activation of the naïve-derived EF pathway, proceeding through aNAV, DN2, and DN3 cells to generate ASC, is characteristic of the patients studied in our cohort, namely, severe Black SLE. Whether this feature also applied to autoreactive B cells and ASC, has remained however, unexplored. Here, as shown in Extended Data Fig. 5a-j, we demonstrate that indeed, the propagation and expansion of Ro60^+^ B cells through an EF pathway is a strong feature of SP patients, but not of HD or the large majority of SN patients. Moreover, we show that this feature is less pronounced in SP patients with LDA.

Combined, our results indicate that HDA in SP patients is characterized by larger accumulations of Ro60^+^ CD27^+^ and CD27^-^ (DN1), switched memory and prominent activation of Ro60^+^ EF cells, in correlation with expansions of ASC. This profile is particularly prominent in about 30% of HDA (cluster C11 in Extended Data Fig. 5e-j), clinically characterized by higher SELENA_SLEDAI. Interestingly, a similar profile was shared by a subcluster of SP patients with LDA in this cross-sectional analysis (cluster C9, 25%), thereby raising the possibility that it could identify impending disease flares. Testing this prediction will necessitate the performance of longitudinal analysis.

Our study extends substantially our understanding of the extent of potentially pathogenic autoreactivity in naïve B cells in HD. In contrast, to previous tolerance studies of global ANA autoreactivity, serologically present in up to 15% of all healthy subjects in the United States or polyreactivity defined through binding to antigens largely unrelated to autoimmune diseases (such as LPS, insulin and ssDNA), we now demonstrate naive B cell autoreactivity against discreet self-antigens associated with SLE and other autoimmune diseases at a similar level in HD and seropositive SLE with a significant degree of polyreactivity against other autoimmune disease antigens in SLE but not in HD. Whether this may be due at least in part, from positive selection promoted by these self-antigens during early B cell development in the SLE bone marrow, remains to be ascertained by future experiments. Nevertheless, these observations provide essential information to understand the pathogenic autoreactivity of naïve-derived plasmablasts we have previously reported in SLE and during the early stages of severe COVID-19 infection^25,36,42^. Moreover, they demonstrate that in contrast to prevailing models postulating that disease-associated human autoantibodies need to be generated through somatic hypermutation, germline-encoded naïve B cells contain a significant degree of autoreactivity of pathogenic potential^58–62^.

Finally, In addition to a better understanding of the regulation of B cell tolerance in HD and SLE, our study provides original evidence regarding ongoing autoimmune reactions in SLE patients with LDA, a state that confers significant protection against flares and accumulation of organ damage ^63^, and which can be achieved after B cell depletion therapy with CART cells or other modalities ^7,64,65^. Nevertheless, we demonstrate that active anti-Ro60 autoimmune reactions leading to the generation of substantial amounts of autoreactive ASC, continue to take place in the majority of SP patients with clinically quiescent disease. This feature, which cannot be established serologically given the stable production of large amount of Ro60 autoantibodies by pre-formed long-lived bone marrow plasma cells, established uncoupling between immunological and clinical remission, and represents a candidate biomarker of future flares and disease outcome. Direct cellular measurements of autoreactivity in residual and repopulating B cells could also represent a more informative assessment of immune resetting and early return of autoimmunity in patients treated with CAR-T cells ^64,66^.

## Materials and Methods

### Healthy and SLE donors

The study was carried out based on approval by Emory University Institutional Review Board (Emory IRB nos. IRB000057983 and IRB000058515), and after adhering to relevant guidelines and regulations. Written informed consent was obtained from each of the donor for collecting blood in heparinized vials and clotting vial for subsequent processing. Healthy donor (n = 27) were recruited in Emory University, and all the SLE samples (n = 223) were recruited from Grady Lupus clinic.

### PBMC isolation and serum/plasma collection

Peripheral blood from heparinized tubes were centrifuged at 500g for 10mins to collect the plasma, and then density gradient centrifugation was carried out at 1000g for 10mins to obtain PBMCs. The PBMCs were washed 3 times in RPMI media, viability was determined by trypan blue staining and live cells were counted using automated hemocytometer. PBMCs were cryopreserved in FBS+10% DMSO for future use. The plasma fraction was further clarified by centrifuging at 2500g for 15mins, and the serum was collected from clotting vials by centrifuging at 800g for 10mins. The plasma and serum fractions were kept frozen in 2-3 aliquots in -80°C freezers.

### Tetramer preparation

For each staining reaction, 80ng of Biotinylated Ro60 (Surmodics) antigen was mixed separately with 85ng of Streptavidin-PE in one tube, and with 75ng of Streptavidin-APC in another tube (6:1 molar ratio), and incubated for 1hour at 4°C. For tetanus specific staining, 100ng of biotinylated tetanus toxoid (Mabtech) was incubated with 50ng of Streptavidin-Pacific Blue in one tube, and with 50ng Streptavidin-AlexaFluor488 in another tube (4:1 molar ratio), and incubated for 1hour at 4°C. The Ro60 and tetanus tetramers freshly prepared were then pooled together in 50μl Brilliant buffer containing 5μM of free D-biotin and Streptavidin-PerCPCy5.5 as decoy probe (antigen-specific cocktail).

### Antigen-specific flow cytometry

PBMCs were thawed and washed with staining buffer, and then stained for surface IgG, followed by Fc receptor blocking. The washed cells were next stained with antigen-specific cocktail (Ro60 or Tetanus) for 1 hour at 4°C. This was followed by staining with surface antibodies for 20mins at 4°C, and then Live/Dead staining at room temperature. Finally, the cells were fixed in 0.8% paraformaldehyde and acquired on Cytek Aurora. The cells were similarly stained for sorting (whenever applicable), only skipping the fixation step.

For intracellular staining, the cells were first stained for Live/Dead, which was followed by Fc blocking and staining with surface antibodies. The cells were them fixed and permeabilized (BD Cytofix/Cytoperm) following manufacturer’s protocol and incubated with anti-IgG and anti-IgA. Following this, the cells were stained with Ro60-antigen specific cocktail for 40mins at 4°C, and then washed with staining buffer and acquired on the same day.

### In-vitro differentiation of switched memory B cells into ASC

Ro60-specific and Ro60 negative switched memory B cells were bulk sorted into sterile 96-well V bottomed plates, at a density of 250 cells per well. These cells were stimulated with IL-2 (50U/ml), IL-21 (10ng/ml), IFN-gamma (20ng/ml), BAFF (10ng/ml) and R848 (1ug/ml) in the presence of 10ug/ml of anti-human IgG and anti-human IgM for 3 days. The media was replaced on day 3 with fresh cytokines and R848 but without anti-IgG and anti-IgM, and cultured for 4 more days. The supernatant was collected at Day7 to be tested by LIPS and ELISA, while the cells were harvested for ELISPOT.

### LIPS assay for Ro60

Ro60-luciferase fusion protein was produced by transfection of Cos1 cells with plasmid Ro60_LUC. 1μl of patient serum was mixed with 1.5x10^5^ light units (LU) of fusion protein in buffer A (50 mMTris, pH 7.5, 100 mM NaCl, 5 mM MgCl2, 1%Triton X-100) and incubated for 1 hour under shaking conditions. The mixture was then transferred to Multi-Screen HTS BV Filter Plates (Millipore Sigma) and incubated for 1 hour at room temperature under shaking with Protein A/G UltraLink® Resin (PierceTM, Thermofisher Scientific). Precipitated auto-antigen luciferase fusion proteins were detected using the Renilla Luciferase Assay System (Promega). Known Ro60 positive patient serum was used to establish a relative unit standard curve and used to determine the relative Ro60 serum titers in the cohort.

### ELISPOT

The wells of MultiScreen-IP sterile filter plates (EMD Millipore) were first washed with 80ul of sterile 70% ethanol and immediately washed 3 times with sterile H20, followed by coating with 20 μg/ml of Ro60 antigen, or tetanus antigen, or 5 μg/ml goat anti-human IgG (Bethyl) in PBS overnight at 4°C. Next day, the plates were washed and blocked with R10 medium for 1hr at 37°C. The cells were seeded at 250 cells/well post in-vitro differentiation in ASC, or as serial dilutions of 1X10^6 PBMCs, and incubated overnight at 37°C. After washing next day, total IgG was detected with 1 μg/ml goat anti-human IgG-alkaline phosphatase (Bethyl) and spots were developed using Vector Blue Alkaline Phosphatase Substrate and then counted using an automated CTL ELISPOT reader.

### Ro60^+^ B cell sorting for single-cell sample preparation

B cell enrichment by was done by Stem Cell B cell enrichment kit (Cat# 19554) from frozen PBMCs of either 1 HD (experiment batch 1) or 4 SLE subjects from SP HDA group (experiment batch 2), then blocked for Fc receptor and stained by Ro60-antigen specific cocktail, followed by cell surface antibody cocktail. For experiment batch 1, healthy donor Ro60^+^ rNAV was directly sorted into 1 tube containing 200ul of RPMI (glutamine-free) + 0.04% Ultrapure BSA. For experiment batch 2, distinct TotalSeq^TM^ hashtag was added for each SLE subject (Biolegend). Finally, cells were stained with live dead stain. B cells from the 4 SLE subjects were pooled together post-hashtagging in FACS buffer (1X PBS + 2% FBS) and anti-Ro60 B cells were sorted in 6 tubes, each containing 200ul of RPMI (glutamine-free) + 0.04% Ultrapure BSA + 4ul of reconstituted TotalSeq™-C Human Universal Cocktail, V1.0 (Biolegend). Ro60⁺ B cells were sorted from enriched parental populations as follows: **pA1** (ASC: CD19⁺ CD27⁺⁺ CD38⁺⁺), **pA2** (enriched aNAV: CD19⁺ IgD⁺ CD21⁻ CD11c⁺), **pA3** (enriched rNAV: CD19⁺ IgD⁺ CD21⁺ CD11c⁻), **pA4** (enriched SwM, DN1, and DN3: CD19⁺ IgD⁻ CD21⁺ CD11c⁻), ^pA5^ (enriched DN2: CD19⁺ IgD⁻ CD21⁻ CD11c⁺), and **pA6** (CD38⁺⁺ B cells) (Extended Data Fig. 4a). Following sorting, cells were washed with RPMI medium and immediately processed for GEM preparation. Each population was loaded separately on a 10x Genomics Chromium controller and processed using the 5’ v2 chemistry to obtain paired gene expression (GEX), surface protein hashtag (HTO), and B-cell V(D)J libraries. Libraries were sequenced on a Novaseq 6000 sequencer and aligned with Cell Ranger multi v7.02 to GRCh38 and the matching VDJ reference. Filtered feature-barcode matrices and the per-sample filtered_contig.fasta and filtered_contig_annotations.csv files were used as input for downstream analysis.

### Gene expression analysis for B cell subset identification

All single-cell analyses were performed in R using DropletUtils, scran, scater, scuttle and batchelor. Gene metadata was retrieved from Ensembl (Jul-2023 archive) through biomaRt. Cells were kept if they had more than 1000 UMIs, more than 750 detected genes, less than 10% mitochondrial reads and less than 40% ribosomal reads. Immunoglobulin related genes were excluded from the gene expression dataset so that immunoglobulin transcripts would not drive the variance, and only protein-coding genes detected with more than 1 count in at least 5 cells were retained. Hashtag identities for subjects were called with and only droplets classified as singlets were kept. Size factors were estimated by deconvolution after a quick clustering step (scran), and counts were log normalized. The top one third of genes by biological variance were used for dimension reduction. The data was integrated across subjects with fastMNN. A shared-nearest-neighbour graph was built on the MNN-corrected components and clustered with the Leiden algorithm. Non-B cell contaminants were removed. B cell subsets were identified from canonical markers as follows: rNAV cells were derived from tube A3 (Ro60⁺ pA3), exhibiting transcriptional signature of high expression of *BACH2, PAX5, FCRL1*, and *CR2*. DN2 cells were derived from tube A5 (Ro60⁺ pA5), showing elevated expression of *ITGAX, TBX21, FCRL5,* and *ZEB2*. Switched memory (SwM) and DN1 cells were both derived from tube A4 (Ro60⁺ pA4). SwM cells were selected based on high expression of *SELL, CD27, CR2*, and *TCF7*, whereas DN1 cells were selected based on high *CXCR5 and TCF7* expression, with absence of *CD27, ITGAX*, and *TBX21* expression.

### B-cell receptor analysis and recombinant monoclonal antibodies

V(D)J contigs from each library were processed with the Immcantation suite of applications. V, D and J gene calls were obtained with IgBLAST using IMGT human germline references. Contigs with the highest UMI count per cell were retained. Only cells passing GEX QC were carried forward. Cells with same V and J genes, junction length and a Hamming distance of 0.15 on the CDR3 nucleotide sequence were assigned as clones. Clonal germlines were reconstructed and somatic hypermutation was scored per sequence. The annotated heavy-chain table was joined back to the SingleCellExperiment object by cell barcode for monoclonal antibody selection.

The VDJ sequences from each subset was selected to span different mutational frequencies and different VH gene family usage. For this study, analyses were restricted to B cell receptor (BCR) sequences to enable monoclonal antibody generation. Negative control monoclonal was made from healthy donor bone marrow resting naïve B cells and shared by Dr. Scott A Jenks. The selected monoclonal antibodies (paired VH-VL sequences) were synthesized by Genscript as IgG antibodies using their Turbo CHO-HT 2.0 expression system.

### Surface plasmon resonance (SPR) for assessing monoclonal binding to autoantigens

SPR was performed and data were collected real-time using benchtop Nicoya Alto SPR platform. Monoclonals (ligands) were captured on to the carboxyl-PEG chip indirectly using Protein A (40ug/ml) by capture kinetics program, and binding of ligands to different analytes were assessed by single-cycle binding kinetics. The regeneration was in Glycine-HCL pH 1.5. The dilutions for each analyte are as follows:

Ro60 (Arotec) and biotinylated tetanus toxoid (Mabtech) were diluted in PBST; biotinylated-Ro60 (Abbexa) was diluted in HBST + 20% glycerol; biotinylated Sm (Arotec) and biotinylated La (Arotec) were diluted in HBST. All the analytes were prepared at a final concentration of 3000nM, and five 3-fold serial analyte dilution was made by the machine.

### Human anti-dsDNA ELISA

96-well high-binding plates (Corning) were coated with Poly-L-Lysine (Sigma) diluted 1:10 in 1X PBS for 1 hour at 37°C. After discarding, the wells were next coated with 5μg/ml of ds-DNA (Human genomic DNA, Roche) in 1X PBS + 0.2% Casein and kept overnight. Next day, plates were blocked with SuperBlock blocking solution (Thermo Fisher) for 1 hour, washed 4 times with PBST and 2 times with PBS; and then the monoclonal antibodies were added at 10μg/ml and incubated for 2hrs. Following washing, AP-conjugated anti-human IgG was added at 1:15000 dilution; plate was developed using substate (1:1 reagent A: reagent B); and the O.D was measured at 650nm.

### Analysis software

The flow cytometry files were analysed by FlowJo V10 software. The graphs and statistical tests were performed using GraphPad Prism software v10.4.2, unless otherwise indicated. Hierarchical clustering was done using the Anthropic Claude AI (Sonnet 4.6) and the online available NG-CHM Heatmap builder developed by MD Anderson Cancer Center. Correlograms were made by packages *corrplot and Hmisc* in R studio.

## Data availability

All the flow datasets and single-cell datasets and analysis code will be available upon request from corresponding authors.

## Acknowledgement

This work was supported by NIH U19AI110483 (I.S), NIH P01AI12580 (I.S. and F.E-H.L.), and Lupus Research Alliance Global Team Science Award (I.S., F.E-H.L., and A.K.). C.C. received funding from Lupus Research Alliance Empowering Lupus Research Career Development Award. We thank Sanz and Lee lab members for their valuable feedback on the study. We thank Dr. Caterina E Faliti for her helpful inputs. We thank Dr. Sung Hoon Cho for helping in proofreading the manuscript. We thank Dr Scott A Jenks for providing the negative control monoclonal antibody. We acknowledge Sanz and Lee lab clinical coordination team for recruiting donors and sample collection. We acknowledge all the healthy donors and SLE patients for participating in this research, as well as the care providers and staff in Grady Hospital and Emory Clinics, Atlanta for cooperating in sample collection process.

## Author Contributions

O.R. and I.S. conceived and directed the study and wrote the manuscript. O.R. processed samples and performed majority of the experiments and data analyses. C.C. performed 10X genomics single-cell RNA sequencing and analyzed the data. C.C. generated figures with R programming and helped with Claude. R.B. and M.G. performed Ro60 LIPS assay. R.B. helped with ELISPOT assay. R.D. helped with monoclonal antibody selection from VDJ data and participated in manuscript review. T.T.P.V. and M.L. helped in sample processing. S.Y.U. and R.T.A identified SLE patient samples for study inclusion and collected samples. A.A.A. and A.K. conducted chart review and scored the patients for disease activity. F.E-H.L. coordinated the collection of samples from healthy donors and provided protocol for ASC generation from memory B cells.

---

**Extended Data Fig. 1.**
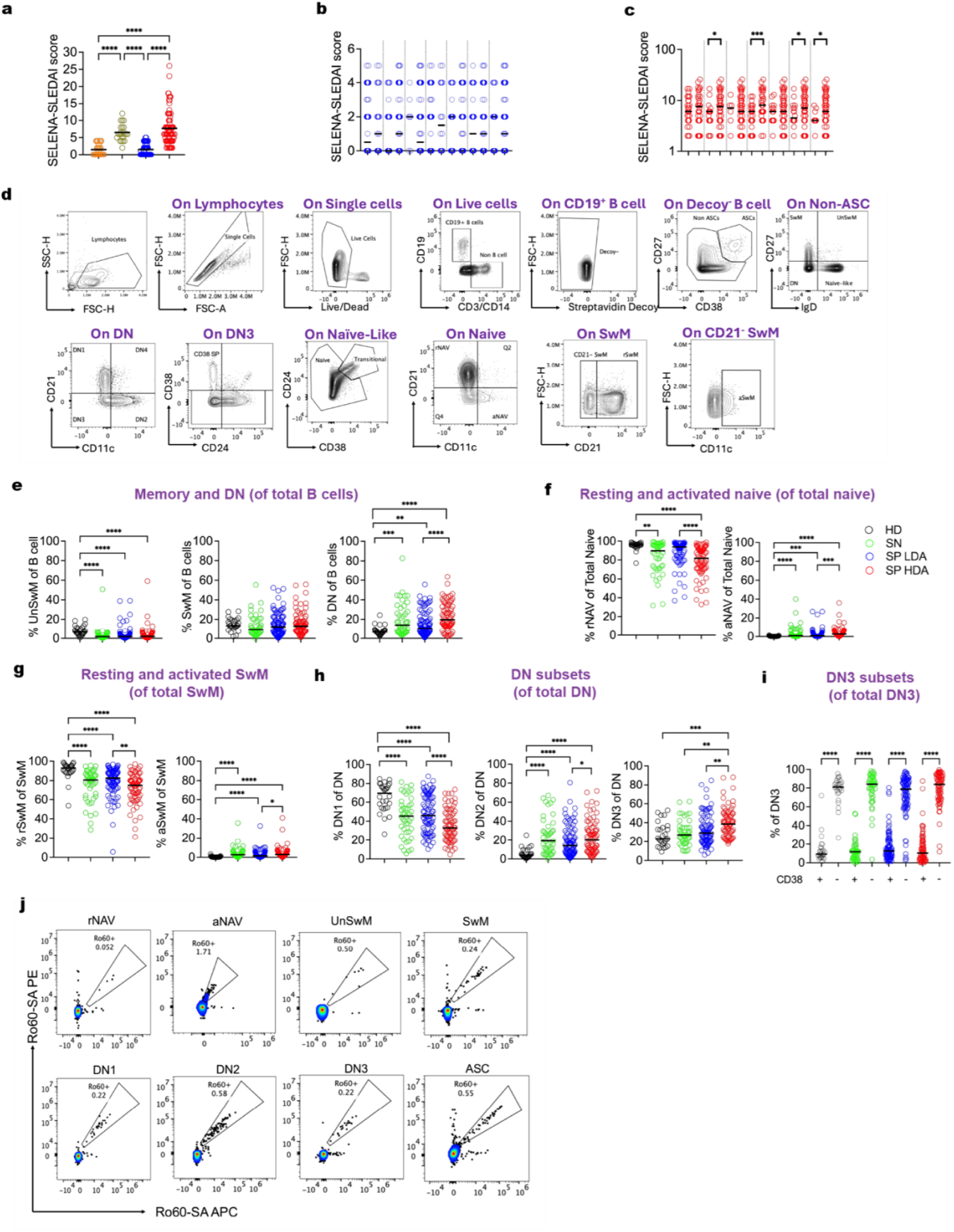
Gating strategy for peripheral B-cell subset immunophenotyping, representative Ro60⁺ flow cytometry profiles across B-cell subsets, and SELENA-SLEDAI scores in SLE patients stratified by presence of Ro60⁺ B cells. **a,** Scatter plots of SELENA_SLEDAI scores of SN and SP patients. **b,c,** Scatter plots of SELENA_SLEDAI scores of LDA SP **(b)** and HDA SP **(c)** patients with (+)/without (-) anti-Ro60 B cells in different subsets (indicated by X-axis). **d,** Representative gating strategy for identifying major B cells subsets in peripheral blood of SLE patient. **e-i,** Scatter plots of frequency of major B cell subsets in different groups. **j,** Representative FACS plots of anti-Ro60 staining in major B cell subsets in SP SLE patient. Horizontal bars in scatter plots represents median, and comparison between median in different groups done by nonparametric Kruskal–Wallis statistical testing using Dunn’s analysis. *P < 0.05; **P < 0.01; ***P < 0.001; ****P < 0.0001.

**Extended Data Fig. 2.**
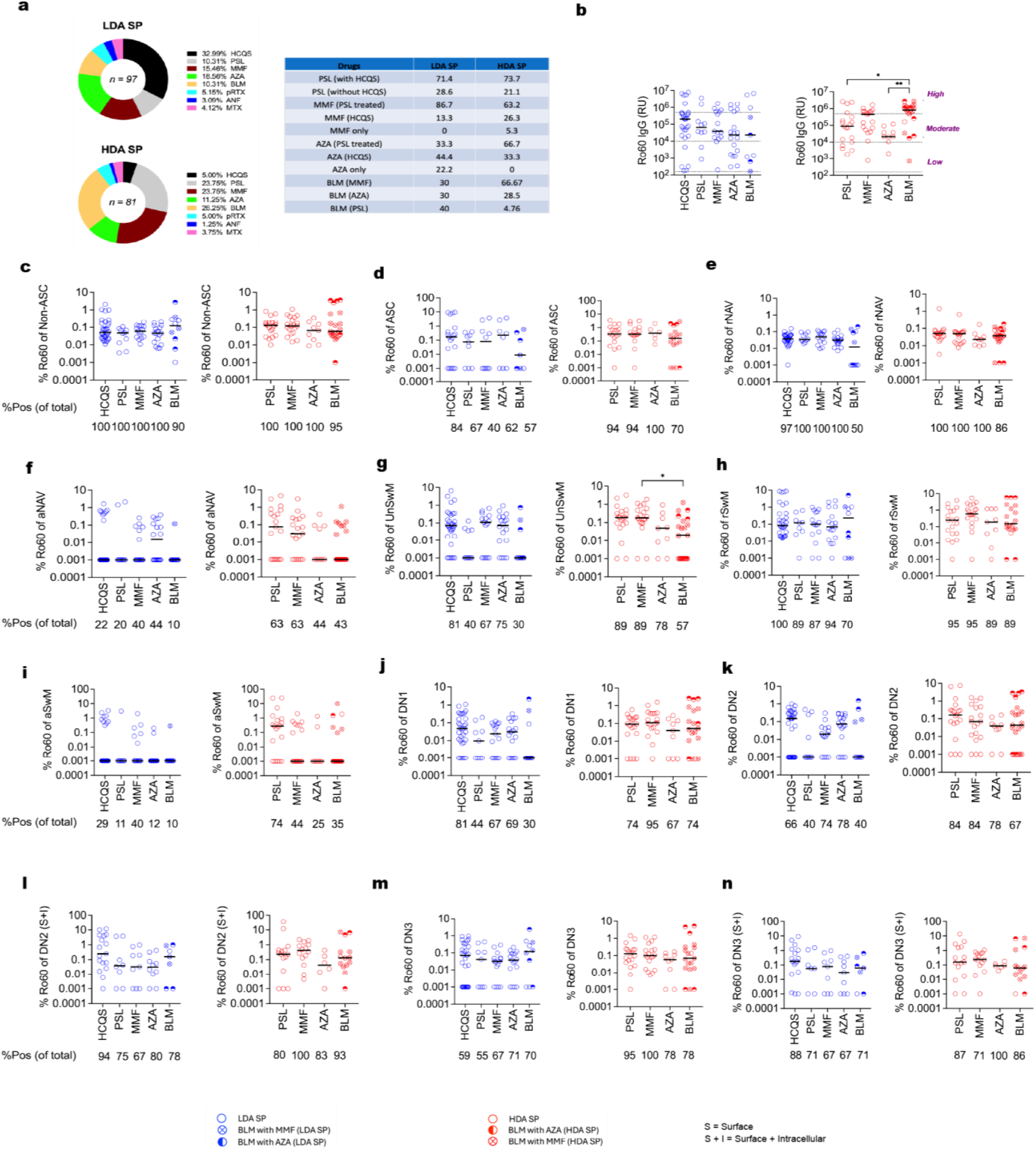
Relationship of treatment regimens on anti-Ro60 B cells frequency in B cell subsets. **a, Left panel:** Donut plot of distribution of different drug regimens in LDA and HDA SP. **Right panel:** Represents the frequency of patients in each drug group (as % in that particular drug group) that received that drug alone or in combination with other drugs. *HCQS: Hydroxychloroquine; PSL: Prednisolone; MMF : Mycophenolate Mofetil; AZA: Azathioprine; BLM: Belimumab; pRTX: post-Rituximab; ANF: Anifrolumab; MTX: Methotrexate*. **b,** Quantification of serum Ro60 IgG by LIPS assay in different treatment groups. **c-n,** Scatter plots of %Ro60 frequency in Non-ASC **(c)**, ASC by ICS **(d)**, rNAV **(e)**, aNAV **(f)**, UnSwM **(g)**, rSwM **(h)**, aSwM **(i)**, DN1 **(j)**, DN2 **(k)**, DN2 by ICS **(l)**, DN3 **(m)** and DN3 by ICS **(n)**. *ICS: Intracellular staining.* All the values are plotted in log scale, values of 0 were replaced by 0.001 so that they can be plotted. Horizontal bars in scatter plots represents median, and comparison between median in different groups done by nonparametric Kruskal-Wallis Kruskal–Wallis statistical testing using Dunn’s analysis. *P < 0.05; **P < 0.01; ***P < 0.001; ****P < 0.0001.

**Extended Data Fig. 3.**
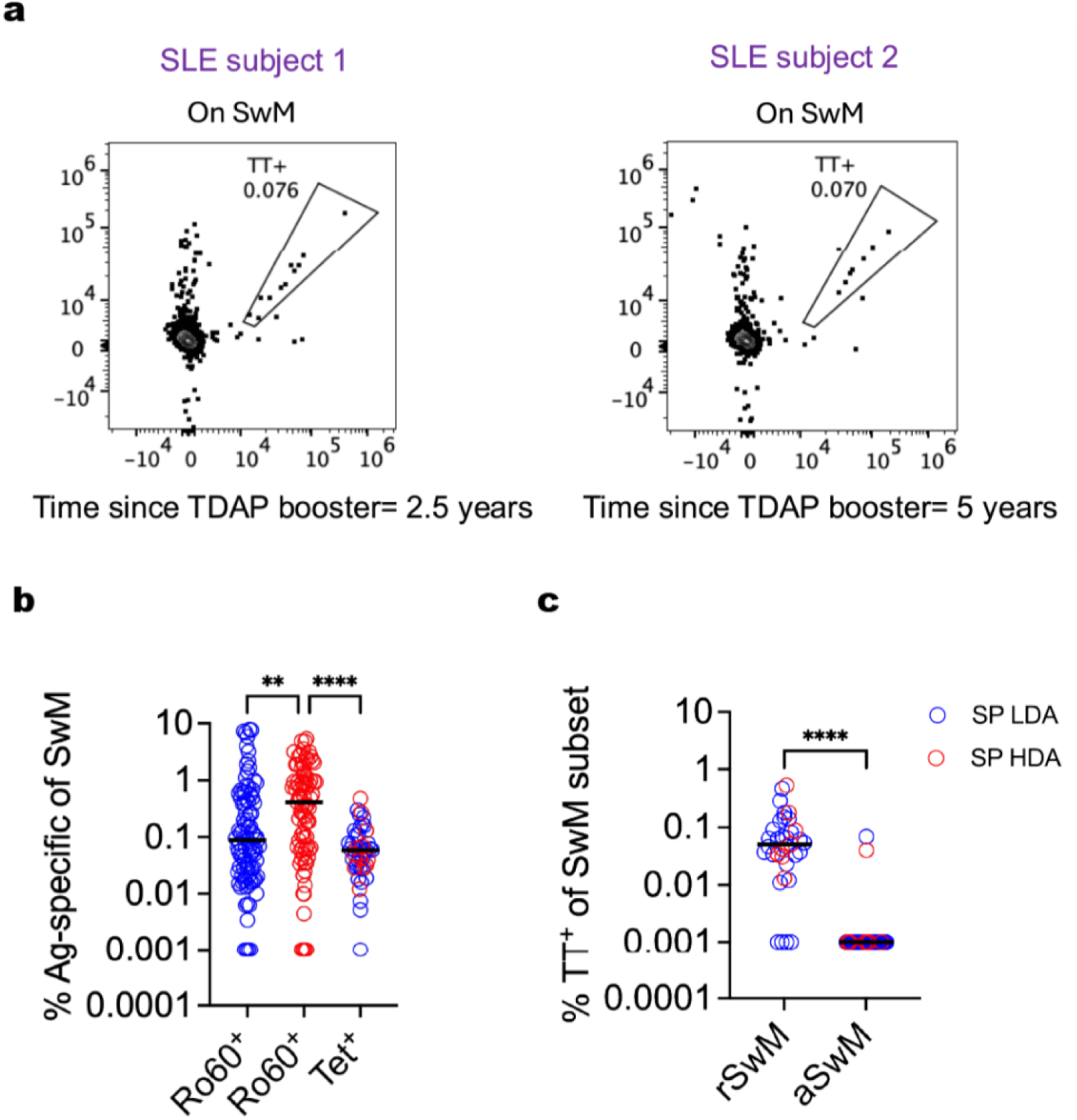
Steady-state tetanus-tetramer staining on SLE patients, comparison of TT^+^ SwM with Ro60^+^ SwM. **a,** Representative FACS plots of steady-state anti-Tetanus staining on SwM in 2 SLE patients, SLE subject 1 had received TDAP booster 2.5 years before draw and SLE subject 2 had received TDAP booster 5 years before draw, as standard of care. **b,** Scatter-dot plot of anti-Ro60 and anti-Tetanus (Tet^+^) SwM in LDA SP(n=34) and HDA SP (n=12) SLE. **c,** Scatter-dot plot of %TT^+^ of rSwM and aSwM in LDA SP(n=34) and HDA SP (n=12) SLE. The patients selected for TT^+^ SwM staining ranged from 4months to 10 years post TDAP booster. All the values are plotted in log scale, values of 0 were replaced by 0.001 so that they can be plotted. Horizontal bars in scatter plots represents median, and comparison between median in different groups done by nonparametric Kruskal-Wallis Kruskal–Wallis statistical testing using Dunn’s analysis. *P < 0.05; **P < 0.01; ***P < 0.001; ****P < 0.0001.

**Extended Data Fig. 4.**
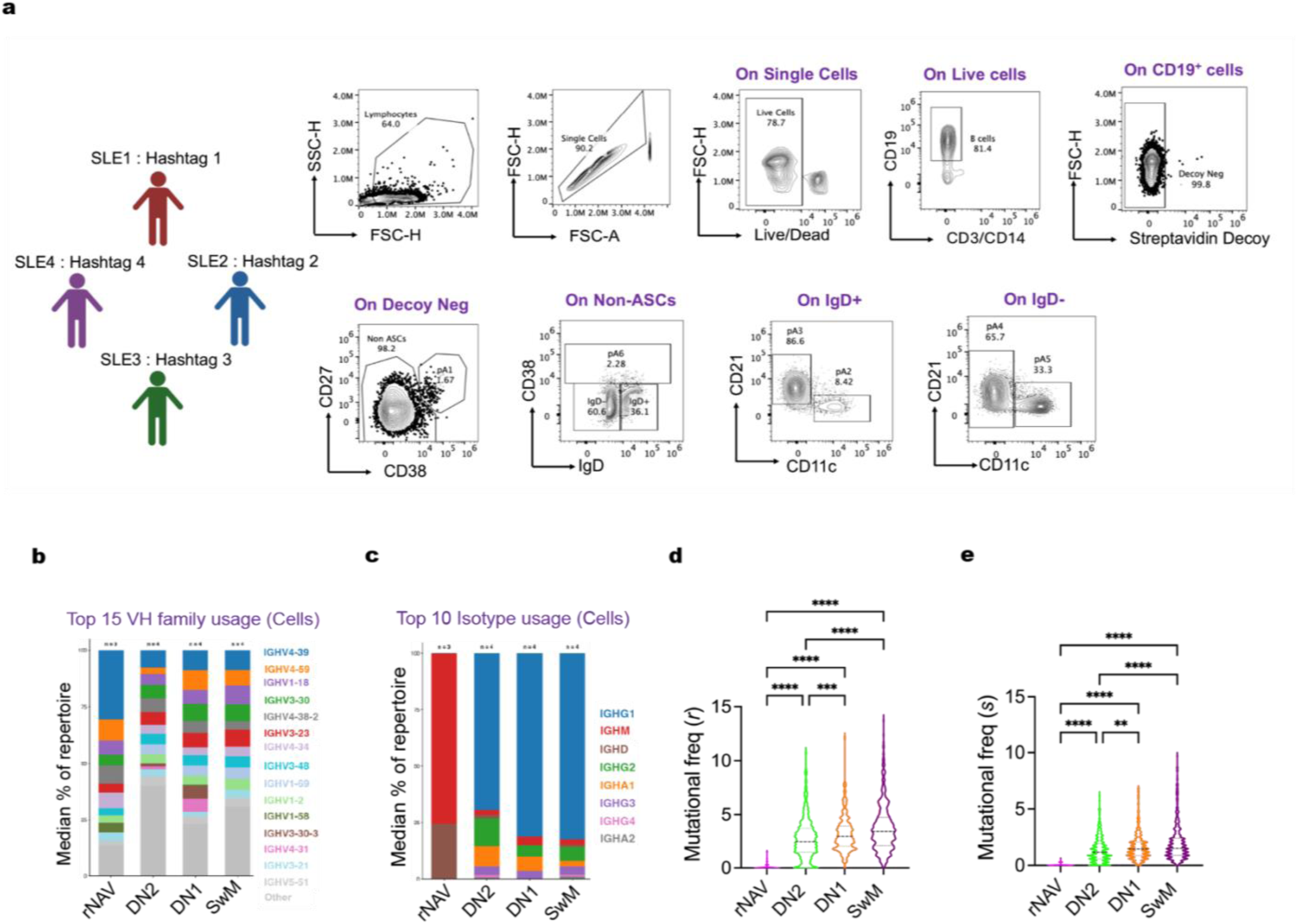
Sorting strategy, VH family usage, isotype usage and mutational frequencies in Ro60^+^ rNAV, DN2, DN1, SwM. **a,** MACS enriched B cells from 4 SLE subject were hash-tagged with 4 different hashtags and Ro60^+^ B cells were sorted from 6 different populations named pA1 (sorted into tube A1), pA2 (sorted into tube A2), pA3 (sorted into tube A3), pA4 (sorted into tube A4), pA5 (sorted into tube A5) and pA6 (sorted into tube A6). **b,c,** Stacked bar-plot for top 15 VH family usage **(b)** and isotype usage **(c)** in sorted single-cells of Ro60^+^ rNAV (#single cell = 48), DN2 (#single cell = 672), DN1 (#single cell = 238), SwM (#single cell = 500). **d, e,** Violin plot of replacement mutational frequencies (denoted by *r*) **(d)** and silent mutational frequencies (denoted by *s*) **(e)** in sorted single cells of Ro60^+^ rNAV, DN2, DN1, SwM. The number of subjects are denoted by *n*. Comparison between different groups done by nonparametric Kruskal–Wallis statistical testing using Dunn’s analysis. *P < 0.05; **P < 0.01; ***P < 0.001; ****P < 0.0001.

**Extended Data Fig. 5.**
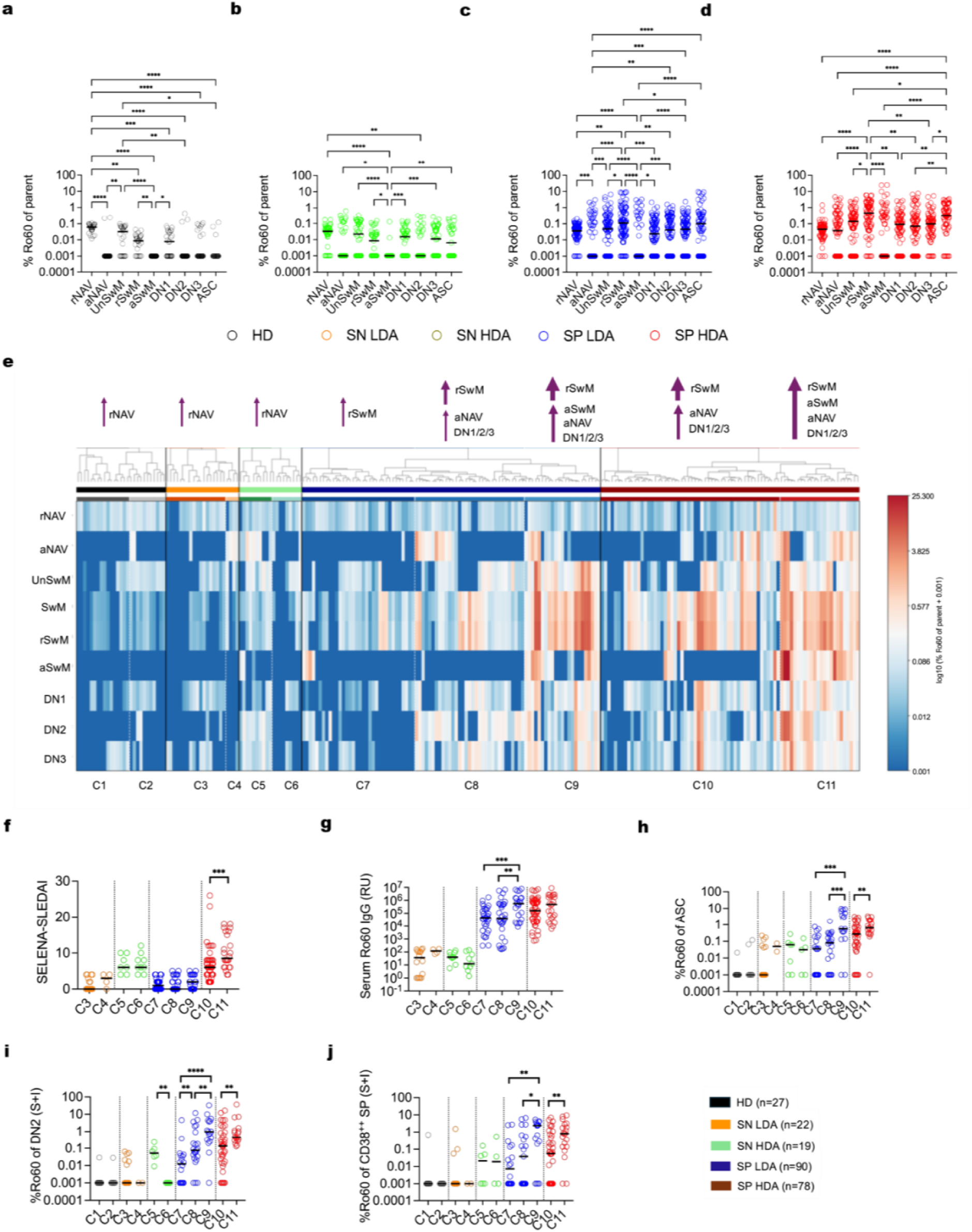
Comparative analysis of Ro60 frequencies across subsets in different groups reveals key features of tolerance of Ro60 autoreactive B cells. **a-d,** Scatter plot of %Ro60 of different B cell subsets in HD **(a)**, SN **(b)**, LDA SP **(c)** and HDA SP **(d)**. **e,** Heatmap of %Ro60 in different subsets, which were categorized into HD, SN LDA, SN HDA, SP LDA and SP HDA. Hierarchical clustering (Ward Linkage) was performed within each group which categorized different groups into different clusters. HD: Cluster C1 (59%), cluster C2 (41%) patients with higher Ro60^+^ UnSwM. SN LDA: Major cluster C3 (82%) and minor cluster C4 (18%) patients with higher Ro60^+^ aNAV. SN HDA: Cluster C5 (53%) patients with higher Ro60^+^ aNAV and DN2 than cluster C6 (47%). SP LDA: Cluster C7 (38%), cluster C8 (37%) patients with higher Ro60^+^ aNAV, DN1/DN2/DN3 than C7; cluster C9 (25%) patients with higher Ro60^+^ aSwM than C7/C8. SP HDA: Cluster C10 (69%), cluster C11 (31%) with more Ro60^+^ aSwM than C10. **f-j)** Scatter plots of SELENA-SLEDAI **(f)**, serum Ro60 IgG **(g)**, %Ro60 of ASC **(h)**, %Ro60 of DN2 by ICS **(i)** and % Ro60 of CD38^++^ SP DN3 by ICS **(j)**, in the different clusters. Horizontal bars in scatter plots represents median, and comparison between median in different groups done by nonparametric Kruskal-Wallis Kruskal–Wallis statistical testing using Dunn’s analysis for **a-c**. Median of clusters were compared within each group by Mann Whitney t-test for plots **f-j**. *P < 0.05; **P < 0.01; ***P < 0.001; ****P < 0.0001.

**Extended Data Fig. 6.**
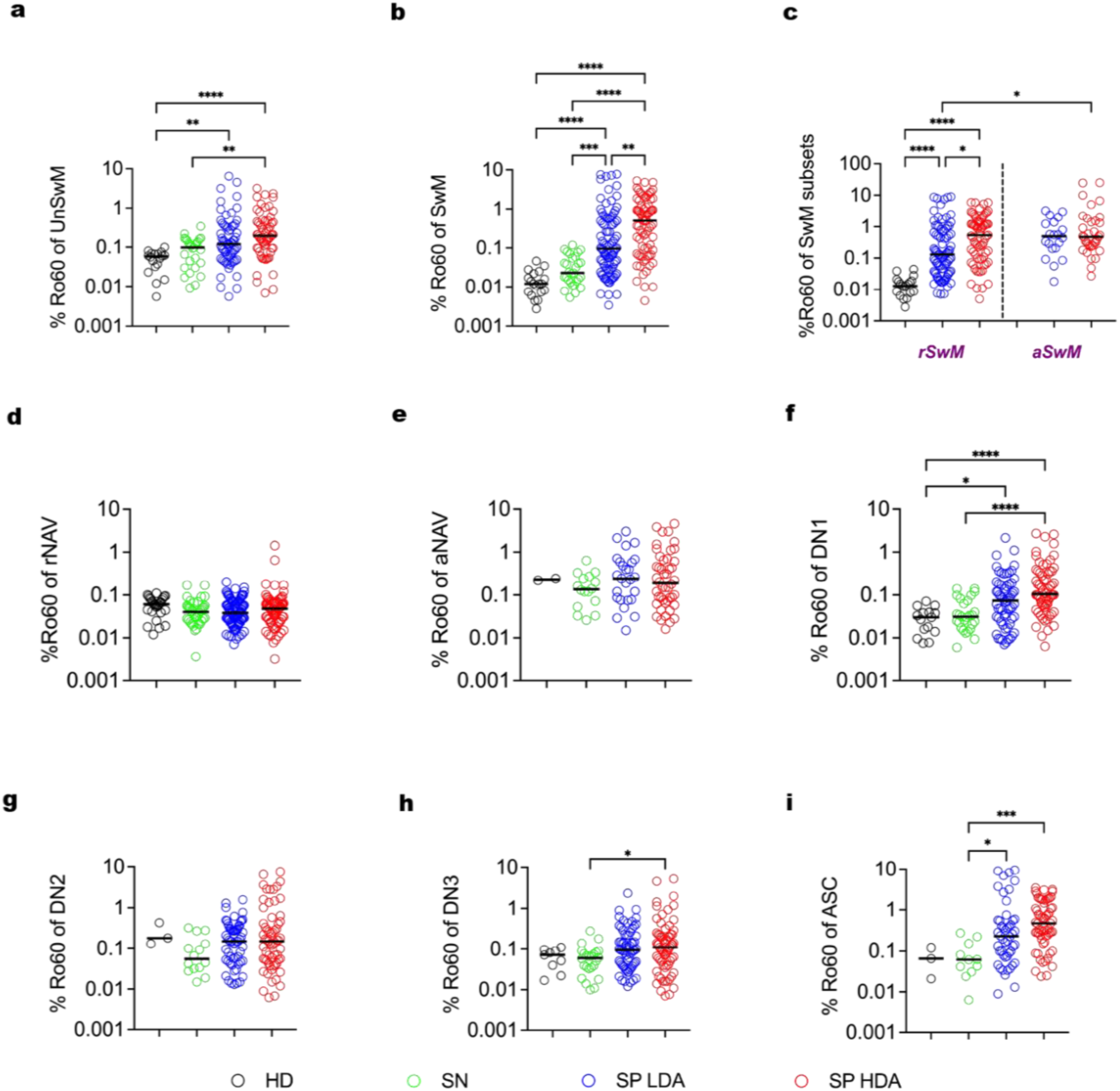
Comparative Analysis of %Ro60⁺ B Cell Subsets Among Groups After Excluding Zeroes. **a-I,** Scatter plots of %Ro60 frequency in UnSwM **(a)**, SwM **(b)** rSwM and aSwM **(c)**, rNAV **(d)**, aNAV **(e)**, DN1 **(f)**, DN2 **(g)**, DN3 **(h)** and ASC **(i)**. All the values are plotted in log scale, values of 0 were replaced by 0.001 so that they can be plotted. Horizontal bars in scatter plots represents median, and comparison between median in different groups done by nonparametric Kruskal-Wallis Kruskal–Wallis statistical testing using Dunn’s analysis. *P < 0.05; **P < 0.01; ***P < 0.001; ****P < 0.0001.

**Extended Data Fig. 7.**
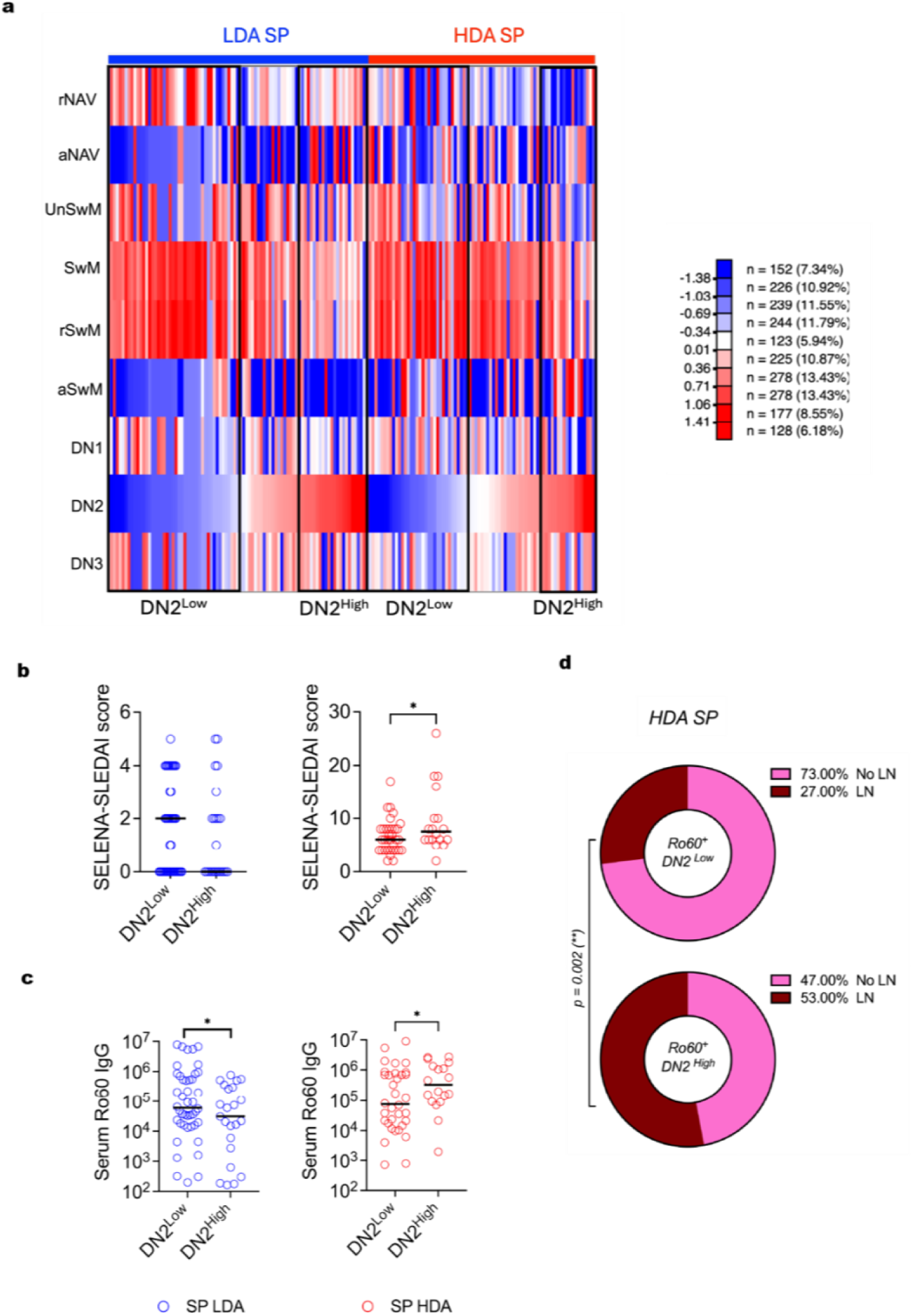
Ro60^+^ DN2 enrichment in HDA SP associates with higher disease activity and prevalent lupus nephritis. **a,** Ro60^+^ B cell frequency in all the subsets for each patient was used to calculate z-scores, and LDA SP and HDA SP patients were then arranged from low to high DN2. A cut-off DN2 z-score of 0.5 was used to identify DN2^High^ patients, while patients with DN2 z-score low than 0.1 were denoted by DN2^Low^. **b,c,** Scatter plots of SELENA_SLEDAI scores **(b)** and serum Ro60 IgG **(c)** in DN2^Low^ and DN2^High^ patients in LDA and HDA SP. **d,** Donut plots of proportion of HDA SP samples which have active LN (lupus nephritis) vs no LN in DN2^Low^ and DN2^High^ patients. Horizontal bars in scatter plots represents median, and comparison between median in different groups done by nonparametric Kruskal-Wallis Kruskal–Wallis statistical testing using Dunn’s analysis. Donut plots were compared by Chi square test. *P < 0.05; **P < 0.01; ***P < 0.001; ****P < 0.0001.

