## Supplementary Information for "Dysregulation of anti-Ro60 B cell autoreactivity in systemic lupus erythematosus"

Table of Contents:

Supplementary Table 1: Antibody panel for Ro60$^+$ B cell flow cytometry

Supplementary Table 2: Patients/healthy donor demographics overview

Supplementary Table 3: SPR $R_{max}$ values and dsDNA ELISA $O.D_{650nm}$ for monoclonal antibodies

Supplementary Table 1: A complete catalogue of antibodies (clone, dilution, cat#) used to classify and sort Ro60$^+$ B cells from different populations

| Antibody Flurochrome | Clone | Dilution | Cat# | Company |
|---|---|---|---|---|
| CD27 BV711 | M-T271 | 1:80 | 356430 | BioLegend |
| CD11c APC-Fire 750 | 3.9 | 1:80 | 301646 | BioLegend |
| CD24 PE-Cy5 | ALB9 | 1:80 | IM2645 | Beckman Coulter |
| IgM BV570 | MHM-88 | 1:80 | 314518 | BioLegend |
| CD3 BUV737 | UCHT1 | 1:160 | 612750 | BD Horizon |
| Streptavidin PerCp Cy5.5 | N/A | 1:200 | 405214 | BioLegend |
| Streptavidin PE | N/A | 1:200 | 405204 | BioLegend |
| CD19 Spark NIR™ 685 | HIB19 | 1:320 | 302270 | BioLegend |
| CD14 BUV737 | M5E2 | 1:320 | 612763 | BD Horizon |
| CD38 BV785 | HIT2 | 1:320 | 303530 | BioLegend |
| CD21 PE-Dazzle 594 | Bu32 | 1:320 | 354922 | BioLegend |
| IgD BV480 | IA6-2 | 1:320 | 566138 | BD Horizon |
| IgG BV421 | M1310G05 | 1:320 | 410704 | BioLegend |
| Streptavidin APC | N/A | 1:500 | SA1005 | Invitrogen |
| CD138 APC -R700 | MI15 | 1:640 | 566050 | BD Horizon |
| Zombie NIR | N/A | 1:1000 | 423106 | BioLegend |

Supplementary Table 2: An overview of demographics of the SLE patients and healthy donors (HD) recruited in the study. **N** represents unique subjects, and **n** represents all samples.

| **Parameters** | **Healthy Donors (HD)** | **Seronegative SLE (SN)** | **LDA Seropositive SLE (SP)** | **HDA Seropositive SLE (SP)** |
|---|---|---|---|---|
| Subjects (**N**) | 27 | 29 | 74 | 50 |
| Samples (**n**) | 27 | 45 | 97 | 81 |
| Age (Mean±S.D) | 37.66 ± 9.8 | 46.27± 14.17 | 45.72± 13.48 | 39.16±11.71 |
| Gender | | | | |
| Female (%) | 92.59 | 100 | 95.16 | 91.89 |
| Male (%) | 7.407 | 0 | 4.83 | 8.1 |
| Race | | | | |
| African-American(%) | 66.67 | 76.67 | 72.58 | 91.9 |
| Caucasian(%) | 14.81 | 13.34 | 9.67 | 2.72 |
| Asian(%) | 18.51 | 3.34 | 1.612 | 0 |
| Not reported (%) | 0 | 6.67 | 8 | 5.45 |

Supplementary Table 3: The SPR $R_{max}$ values and dsDNA ELISA $O.D_{650nm}$ (Blank Subtracted) for monoclonal antibodies have been indicated. For SPR, any Rmax values less than 20 has been denoted by (-) and indicated no measurable binding.

| mAB code | Unbiot -Ro60 $RU_{max}$ | Biot Ro60-SA $RU_{max}$ | Biot La-SA $RU_{max}$ | Biot Sm-SA $RU_{max}$ | Biot Tet-SA $RU_{max}$ | ds DNA ELISA ($O.D_{650nm}$) (Blank subtracted) |
|---|---|---|---|---|---|---|
| Neg C mAb | - | - | - | - | - | 0.026 |
| HDRN1 | - | 423 | - | - | - | 0.048 |
| HDRN2 | - | 618 | 186 | - | - | 0 |
| HDRN3 | 117 | 605 | - | - | - | 0.011 |
| HDRN4 | - | - | - | - | - | 0.011 |
| LRN1 | - | 406 | 790 | 324 | - | 0.088 |
| LRN2 | - | 502 | 1339 | 719 | - | 0.016 |
| LRN3 | - | - | 530 | - | - | 0.033 |
| LRN4 | - | 743 | 348 | 35 | - | 0.002 |
| LRN5 | - | - | - | - | - | -0.004 |
| LRN6 | - | - | - | - | - | 0.011 |
| LRN7 | - | - | - | - | - | 0.006 |
| LRN8 | - | 249 | - | - | - | 0.004 |
| LRN9 | - | 574 | - | - | - | 0.003 |
| LRN10 | - | 59 | - | 262 | - | -0.002 |
| LRN11 | - | 755 | 179 | - | - | 0.215 |
| LRN12 | - | - | - | - | - | 1.011 |
| LRN13 | - | 1117 | 129 | - | - | 0.113 |
| LRN14 | 100 | 887 | - | - | - | 0.024 |
| LRN15 | - | 36 | - | - | - | -0.001 |
| LDN1 | 625 | 1215 | - | - | - | -0.009 |
| LDN2 | - | 40 | - | - | - | -0.006 |
| LDN3 | - | 270 | 700 | - | - | 0.015 |
| LDN4 | - | - | - | - | - | 0.006 |
| LDN5 | - | 115 | - | 563 | - | 0.036 |
| LDN6 | - | 405 | 233 | 467 | - | 0.096 |
| LDN7 | - | 289 | 155 | - | - | 0 |
| LDN8 | 345 | 530 | - | - | - | 0.019 |
| LDN9 | - | - | - | - | - | -0.009 |

| | | | | | | |
|---|---|---|---|---|---|---|
| LDN10 | - | 392 | - | - | - | 0.004 |
| LDN11 | - | 160 | - | - | - | 0.136 |
| LDN12 | - | - | - | - | - | 0.031 |
| LDN13 | - | 342 | - | 469 | - | 0.017 |
| LDN14 | 51 | 74 | - | - | - | 0.055 |
| LDN15 | - | - | - | - | - | 0.058 |
| LDN16 | - | 391 | - | - | - | 0.051 |
| LSW1 | 642 | 1167 | - | - | - | -0.009 |
| LSW2 | 502 | 1082 | - | - | - | -0.01 |
| LSW3 | 240 | 408 | - | - | - | -0.002 |
| LSW4 | 336 | 784 | 64 | - | - | -0.005 |
| LSW5 | 737 | 890 | - | - | - | 0.005 |
| LSW6 | - | 57 | - | - | - | 0.035 |
| LSW7 | - | - | - | - | - | 0.015 |
| LSW8 | 167 | 220 | - | 459 | - | 0.001 |
| LSW9 | - | 230 | - | - | - | 0.095 |
| LSW10 | - | - | - | - | - | 0.002 |
| LSW11 | 625 | 855 | - | - | - | 0.012 |
| LSW12 | 455 | 149 | - | - | - | 0.04 |
| LSW13 | - | 116 | - | - | - | 0.012 |
| LSW14 | 566 | 815 | - | - | - | 0.076 |
| LSW15 | - | - | - | - | - | 0.168 |
| LDN1 1 | - | - | - | - | - | 0.009 |
| LDN1 2 | 122 | 1134 | 920 | 913 | - | -0.01 |
| LDN1 3 | 588 | 1138 | 512 | 613 | - | 0.041 |
| LDN1 4 | 662 | 1113 | - | - | - | -0.012 |
| LDN1 5 | - | - | - | - | - | -0.005 |
| LDN1 6 | - | 456 | - | - | - | 0.02 |
| LDN1 7 | - | - | - | - | - | -0.007 |
| LDN1 8 | - | 1095 | 516 | - | - | -0.003 |
| LDN1 9 | 657 | 199 | - | - | - | 0.005 |
| LDN1 10 | - | - | - | - | - | -0.005 |
| LDN1 11 | - | 183 | - | 570 | - | -0.011 |
| LDN1 12 | 645 | 1050 | - | - | - | -0.009 |
| LDN1 13 | - | - | - | - | - | -0.008 |
| LDN1 14 | 542 | 908 | 80 | - | - | 0 |
| LDN1 15 | - | - | - | - | - | 0.019 |
